# Spatially-resolved single cell atlas of liposarcoma reveals lineage hierarchies, immune niches, and regulatory circuits

**DOI:** 10.64898/2026.03.23.713651

**Authors:** Ryan A. Denu, Veena Kochat, Zhao Zheng, Suresh Satpati, Danh D. Truong, Emre Arslan, Corey Weistuch, Margarita Divenko, Manrong Wu, William Padron, Davis R. Ingram, Khalida M. Wani, Wei-Lien Wang, Sharon M Landers, Hannah C. Beird, Jamie L. McCuisto, Alysha Simmons, Liz Marie Albertorio-Sáez, Danielle N. Maryanski, Cheryl C. Szany, Bryan J. Venters, Carolina Lin Windham, Michael-Christopher Keogh, Keila E. Torres, Christina L. Roland, Emily Z. Keung, Elise F. Nassif Haddad, Alexander J. Lazar, Joseph A. Ludwig, Neeta Somaiah, Kunal Rai

## Abstract

Well-differentiated and dedifferentiated liposarcoma (WDLPS and DDLPS) exhibit markedly different clinical behaviors, with DDLPS showing greater aggressiveness, higher recurrence and metastasis rates, and worse outcomes. Using single-nucleus multiome sequencing, epigenomic profiling, and spatial transcriptomics, we characterized cellular and epigenetic heterogeneity between these subtypes at single-cell and spatial resolution. We found distinct phenotypic states reflecting altered lineage differentiation and plasticity: DDLPS is dominated by early-differentiated progenitor-like cells, sclerotic WDLPS displays broader mesenchymal lineage plasticity, and adipocytic WDLPS contains abundant committed adipocytes. The DDLPS immune microenvironment was dominated by immunosuppressive macrophages, whereas WDLPS harbored more T cells and inflammatory macrophages. Notably, sclerotic WDLPS displayed intermediate cellular and molecular features, suggesting it may represent a distinct WDLPS subtype. Importantly, we identified novel gene regulatory circuits underlying each state, including FABP4/PPARG programs in adipocytic WDLPS, GLI2/TCF7L2/RBPJ/KLF7 programs in sclerotic WDLPS, and KLF7/FOSL2/SP3/GLI2/RBPJ programs in DDLPS. H3K27ac-marked enhancers were enriched near adipocytic marker genes in WDLPS and mesenchymal markers in DDLPS. Together, these findings reveal the cellular heterogeneity of tumor and immune compartments across liposarcoma subtypes and identify regulatory programs driving their differentiation states.

**GRAPHICAL ABSTRACT:** 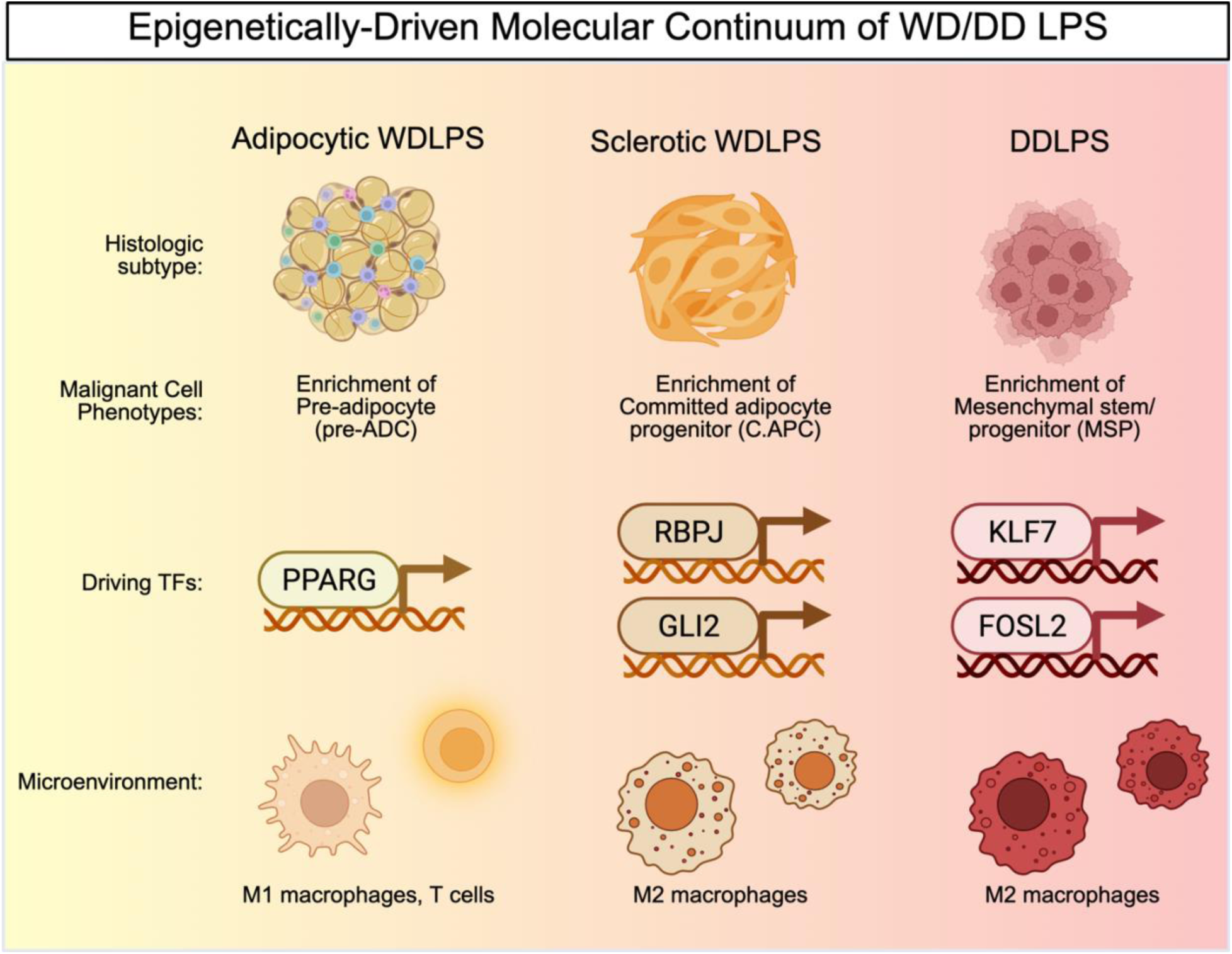

## INTRODUCTION

Liposarcoma (LPS) is the most common form of soft tissue sarcomas, accounting for 20% of all sarcomas, and can arise in any anatomic site, most commonly in the retroperitoneum, extremities, and trunk.^1–3^ LPS is classified into three distinct types with significantly different biology and clinical behavior based on morphology and cytogenetics: well-differentiated/dedifferentiated (WD/DD LPS), characterized by chromosome 12q13-15 amplification, myxoid (characterized by the *FUS-DDIT3* fusion), and pleomorphic (complex genome).^4–6^ WD/DD LPS is the most common of these three LPS subtypes. On one end of this disease spectrum, WDLPS is a low-grade adipocytic tumor thought to arise from mature adipocytes and has a high risk of local recurrence after surgery.^7^ About 10-20% of WDLPS will progress to DDLPS, a more aggressive form of adipocytic tumors, which frequently recurs locally and has a propensity for distant metastasis in 15-30% of cases.^8–12^ More often, DDLPS presents *de novo* at the time of diagnosis rather than arising from pre-existing WDLPS.^13,14^ WDLPS is further divided into two histological subtypes of sclerotic and adipocytic (also lipoma-like or minimally sclerotic) LPS. Other terms have been used to describe differences in this spectrum of disease, including cellular WDLPS and low-grade DDLPS.^10,15^ Adipocytic WDLPS has been associated with improved recurrence-free and overall survival compared to sclerotic WDLPS.^16^ Effective treatments for unresectable, recurrent, or metastatic disease are lacking and remain a critical unmet need for this disease.

Previous studies have attempted to identify the factors that promote the morphogenesis from WDLPS to DDLPS. Initial studies suggested that DDLPS harbors about twice as many chromosomal copy aberrations, including recurrent amplifications in 19q13.2, *GLI1*, *JUN*, and *MAP3K5*.^14,17^ Genomic profiling of matched WDLPS and DDLPS from 17 patients revealed that only 8.3% of somatic mutations in WDLPS were shared with DDLPS.^18^ Further, DDLPS showed more chromosome 12q amplifications, copy number losses, and gene fusions, particularly those involving differentiation genes *HMGA2* and *CPM*.^18,19^ However, some studies failed to detect genetic differences between WDLPS and DDLPS.^20^

Our previous work^21^ suggested that epigenetic reprogramming may contribute to the development of DDLPS. Compared with WDLPS, DDLPS was associated with higher levels of the repressive histone mark H3K9me3, higher levels of the active enhancer mark H3K27Ac, lower levels of the active transcription mark H3K79me3, and lower levels of 5-hydroxymethylcytosine.^21^ Functionally, H3K9me3-induced epigenetic reprogramming, such as that driven by KLF6, plays a key role in dedifferentiation.^21,22^ Recent single-cell studies have begun to characterize the heterogeneity of cellular populations within WDLPS and DDLPS.^23,24^

We hypothesized that the extensive cellular diversity observed in LPS arises from intrinsic cellular plasticity and metastable epigenetic states that are susceptible to reprogramming, enabling transformation between WDLPS and DDLPS states. We provide comprehensive maps of LPS at the transcriptomic, epigenetic, and spatial levels using complementary multi-omics technologies. First, we analyzed twenty-three LPS samples using snRNA-seq and snATAC-seq. Next, we performed spatial transcriptomics on three paired samples, revealing a remarkable degree of cellular, epigenetic, and spatial heterogeneity. Finally, we applied non-negative matrix factorization and core regulatory circuitry analysis to identify transcriptional regulatory programs that drive specific cellular phenotypes and define novel spatial niches. Our findings suggest that sclerotic WDLPS harbors cellular and epigenetic phenotypes that are more similar to DDLPS than to adipocytic WDLPS.

## RESULTS

### Gene expression and epigenome atlas of LPS dissects underlying cellular diversity and molecular heterogeneity

The intratumoral heterogeneity and cellular diversity of WD/DD LPS and the epigenetic programs underlying aberrant differentiation remain poorly defined. To address this, we performed snMultiome sequencing (simultaneous snRNA-seq and snATAC-seq) on WDLPS and DDLPS tumors from 13 treatment-naïve patients (23 samples total: 15 WDLPS from 10 patients and 8 DDLPS from 5 patients; **Supplementary Tables S1–S2**). Among WDLPS tumors, 5 were adipocytic and 10 sclerotic according to established criteria. Nine tumors (69.2%) were retroperitoneal, 11 (84.6%) were stage IIIB, and the mean tumor size was 20.8 cm (**Supplementary Table S3**). Three specimens contained matched WDLPS and DDLPS components dissected from the same tumor (**Figure 1A; Supplementary Figure S1**). We obtained high-quality profiles from 90,553 and 90,395 nuclei by snRNA-seq and snATAC-seq, respectively. snRNA-seq analysis identified 44 clusters, annotated using differentially expressed genes to define cellular identities and states (**Figure 1B**, **Supplementary Figure S2A**). Tumor cells were distinguished from non-malignant populations using complementary approaches. InferCNV was used to identify gains in 12q13, which includes the hallmark amplification of *MDM2* (**Figure 1C, Supplementary Figure S2B and S2C**). CytoTRACE scores were used to determine stemness, which were anticipated to be higher in malignant cells (**Figure 1D, Supplementary Figure S3**). UMAP embedding and cluster-specific gene expression further resolved major cell types, including tumor cells (*CDK4*, *MDM2*), macrophages (*CD163*, *ITGAM*, *CD74*), T cells (*CD4*, *CD8A*, *CD3D*, *GZMA*), endothelial cells (*PECAM1*, *SNTG2*, *VWF*), dendritic cells (*CLEC4C*, *CD1C*), myocytes (*MYOG*, *TRDN*), smooth muscle cells (*TRPC6*, *NFASC*, *MYH11*), and mast cells (*CPA3*, *SLC18A2*) (**Figure 1E; Supplementary Table S4**).

**Figure 1.**
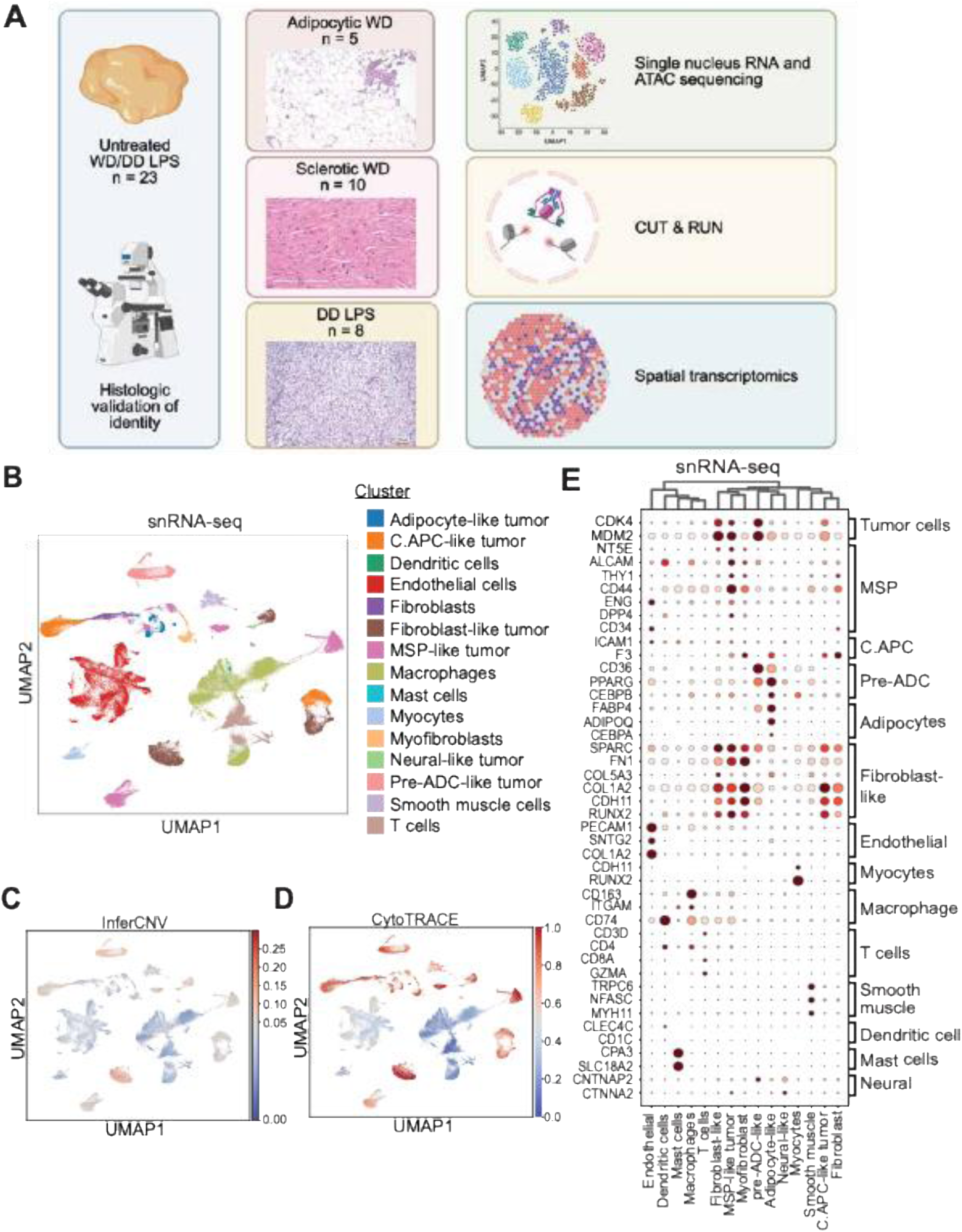
Single cell characterization of LPS reveals cellular diversity and molecular heterogeneity. (A) Schematic overview of the experimental workflow. Resected LPS were dissected by a pathologist and classified as WDLPS (adipocytic vs sclerotic) or DDLPS. Single nucleus multiome was then performed, followed by integration of snRNA-seq and snATAC-seq data and downstream analysis. (B) UMAP analysis of snRNA-seq showing annotations by cell type. (C) Heatmap of inferCNV scores projected onto snRNA-seq UMAP. (D) Heatmap of CytoTRACE scores projected onto snRNA-seq UMAP. (E) Heat map showing expression of differentially expressed genes in each cluster.

### LPS cells exhibit a spectrum of differentiation states reflecting mesenchymal lineage plasticity

A key determinant of clinical classification in LPS is tumor cell differentiation status, which pathologists use to distinguish WDLPS from DDLPS. We therefore defined tumor cell states using three complementary criteria: established lineage markers, overlap with transcriptomic profiles from a mesenchymal stem cell (MSC) differentiation model, and CytoTRACE-derived differentiation scores. Malignant cell clusters were further subdivided according to adipocyte differentiation markers: mesenchymal stem/progenitor (MSP)-like tumor cells (*NT5E, ALCAM, THY1, CD44, ENG, DPP4, CD34*); fibroblast-like tumor cells (*SPARC, FN1, COL5A3, CDH11, RUNX2*); committed adipose progenitor-like tumor cells (C.APC; *ICAM1 and F3*); pre-adipocyte-like tumor cells (*pre-ADC; CD36, PPARG, CEBPB);* and adipocyte-like tumor cells (*FABP4*, *ADIPOQ*, *CEBPA*) (**Figures 1B and 1E, Supplementary Figure S4**).^25–28^ Interestingly, we also identified a smaller population of neural-like tumor cells expressing *CNTNAP2* and *CTNNA2*.

To define the differentiation hierarchy of LPS cells, we leveraged an *in vitro* human MSC differentiation model comprising twelve transcriptional archetypes representing distinct lineage states (**Figure 2A**).^29^ These included undifferentiated mesenchymal (archetypes 1–2), mesenchymal progenitor (3–4), adipocytic (5–6), osteogenic (7), and chondrogenic states (8–12). Tumor clusters identified by snRNA-seq (**Figure 1B**) were projected onto this framework to infer their differentiation states (**Figure 2B**). Adipocytic WDLPS tumors were strongly enriched for adipocytic archetype 6, consistent with a more differentiated phenotype resembling normal adipocytes. In contrast, DDLPS samples were enriched for progenitor-like archetypes 3 and 4. Sclerotic WDLPS and DDLPS tumors showed enrichment of chondrogenic archetype 12, whereas osteogenic archetype 7 was specifically associated with sclerotic WDLPS. Notably, undifferentiated archetype 1 was enriched in MSP-like clusters, with highest expression in DDLPS and sclerotic WDLPS, indicating a more primitive mesenchymal state. Archetype 5, representing an intermediate adipocytic state, was broadly represented across LPS subtypes.

**Figure 2.**
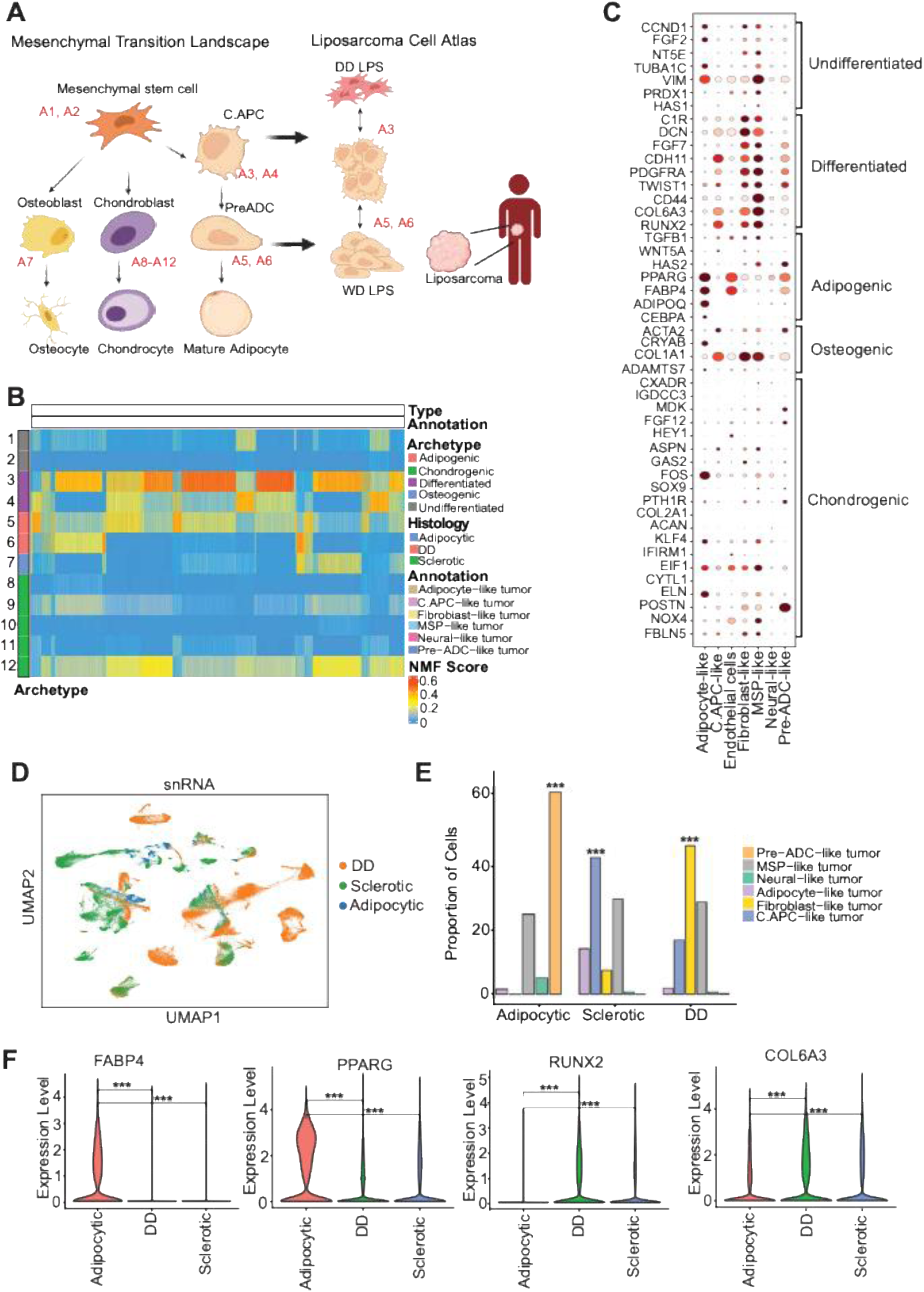
Dissecting WD and DD LPS clusters based on lineage-specific archetypes. (A) Lineage-specific archetype data from an *in vitro* MSC differentiation model were integrated with snRNA and snATAC data from WD/DD LPS to identify the differentiation states of tumor cells. (B) Lineage-specific archetype score heatmap of WD and DD LPS patient samples. Row annotations indicate the lineages specified by each archetype. (C) Bubble plots showing the expression of representative markers of each archetype by snRNA-seq. (D) UMAP of snRNA-seq data showing differences based on sample identity: adipocytic WD, sclerotic WD, or DD. (E) Proportion plot showing distribution of tumor cell types in adipocytic WD, sclerotic WD, or DD. (F) Comparison of expression of mesenchymal markers (*RUNX2* and *CDH11*) and adipocytic markers (*ADIPOQ* and *PPARG*) between WD (adipocytic and sclerotic) and DD LPS components. *** P < 0.0001.

To define dominant transcriptional programs across tumor cells, we performed archetype analysis on snRNA-seq tumor clusters. Each cluster exhibited a distinct combination of archetypes, revealing both shared and cluster-specific transcriptional states (**Supplementary Figure S5A**). UMAP embedding of archetypes recapitulated the organization of snRNA-seq clusters, indicating that archetypes capture biologically meaningful tumor cell programs (**Supplementary Figure S5B**). Visualization of archetype intensity highlighted spatial gradients for selected archetypes, including archetypes 3, 6, and 7, across the tumor cell manifold (**Supplementary Figure S5C**). Quantitative comparison of archetype intensities across clusters further confirmed differential enrichment of specific archetypes within defined tumor populations (**Supplementary Figure S5D**). For each archetype, we identified genes whose expression most strongly correlated with archetype intensity, defining distinct transcriptional signatures underlying each program (**Supplementary Figures S5E–P**). Collectively, mapping LPS cells onto a normal differentiation landscape spanning MSCs to mature adipocytes indicates that DDLPS tumors persist in a progenitor- or stem-like state. In contrast, sclerotic WDLPS tumors exhibit broader mesenchymal lineage plasticity, with partial osteogenic and chondrogenic features.

To further interrogate lineage states, we examined expression of differentiation-associated markers. Adipocyte- and pre-ADC-like tumor clusters expressed canonical adipocyte genes (*HAS2, PPARG, FABP4, ADIPOQ, CEBPA*; **Figure 2C**). In contrast, less differentiated clusters, including C.APC-, MSP-, and fibroblast-like tumor cells expressed mesenchymal and stromal lineage markers (*DCN, FGF7, CDH11, PDGFRA, TWIST1, CD44, COL6A3, RUNX2*), with MSP clusters expressing additional undifferentiated MSC markers (*THY1, NT5E, PRDX1*). These lineage-specific signatures further position LPS subtypes along an MSC-to-adipocyte differentiation continuum.

We next examined the distribution of tumor cell states across histologic subtypes of liposarcoma (**Figure 2D**). Adipocytic WDLPS tumors were enriched for differentiated pre-ADC-like tumor cells, whereas sclerotic WDLPS and DDLPS tumors showed increased representation of less differentiated MSP-like states (**Figure 2E**). C.APC-like tumor cells were enriched in sclerotic WDLPS, while fibroblast-like tumor cells were enriched in DDLPS. Consistently, adipocytic genes *FABP4* and *PPARG* were highly expressed in adipocytic WDLPS, whereas fibroblastic genes *RUNX2* and *COL6A3* were upregulated in sclerotic WDLPS and DDLPS (**Figure 2F**).

Individual tumor clusters also displayed strong subtype bias, with distinct proportions of cells originating from adipocytic WDLPS, sclerotic WDLPS, or DDLPS tumors, indicating subtype-specific transcriptional programs (**Supplementary Figure S6A**). Similar trends were observed when tumor clusters were analyzed in aggregated groups, confirming robust subtype-associated distributions (**Supplementary Figure S6B**). Absolute cell counts further highlighted differences in the abundance of tumor cell clusters across subtypes (**Supplementary Figure S6C**), while tumor-level analyses revealed substantial inter-tumoral heterogeneity within each subtype (**Supplementary Figure S6D**). In contrast, benign cell populations displayed more conserved distributions across subtypes, although immune cells were enriched in DDLPS (**Supplementary Figure S6E**).

### Patient-wise heterogeneity in de-differentiation trajectory and cell states during LPS progression

To directly compare tumor cell states between WDLPS and DDLPS, we analyzed snRNA-seq data from three matched synchronous WD/DD LPS tumor pairs. Joint UMAP embedding showed intermixing of cells from paired tumors while preserving tumor-specific structure (**Supplementary Figure S7A**). Unsupervised clustering identified 36 transcriptionally distinct cell populations, annotated using canonical marker gene expression (**Supplementary Figures S7B–C**). Mapping *MDM2* expression enabled clear identification of malignant WD/DD tumor cells, consistent with *MDM2* amplification as a defining genomic feature of these tumors (**Supplementary Figure S7D**). Comparative analysis revealed systematic shifts in the relative abundance of cell populations between WDLPS and DDLPS tumors, with PreADC-like tumor cells preferentially enriched in WDLPS samples and fibroblasts-like tumor cells enriched in DDLPS samples (**Supplementary Figure S7E**).

We next examined the differentiation trajectories within these three matched WD/DD LPS tumor pairs (**Figure 3A**). Analysis of cluster differences between the WDLPS and DDLPS components of these tumors demonstrated enrichment of pre-ADC-like tumor cells in WDLPS and fibroblast-like tumor cells in DDLPS (**Figure 3B**). Normalized non-negative matrix factorization (N-NMF) archetype analysis further demonstrated enrichment of undifferentiated or more mesenchymal stem-like archetypes (e.g. archetype 3) in DDLPS as compared to its matched adipocytic WDLPS, while its matched adipocytic WDLPS component demonstrated enrichment of more adipocytic archetypes (e.g. archetypes 5-6). However, for patient 10, there are less obvious differences between the WDLPS and DDLPS components (**Figure 3C**). Trajectory analysis of individual tumor pairs revealed distinct differentiation structures (**Figure 3D**). In patient 1, whose WDLPS component was adipocytic in nature, we noted a number of nodes and branches of differentiation. On the other hand, tumors from patients 4 and 10, where the WDLPS component is sclerotic in nature, showed fewer major nodes and branches of differentiation. Assuming the most differentiated cell type approximates the cell-of-origin, these analyses suggest that adipocytic WDLPS and DDLPS occupy more distant positions in a differentiation continuum, whereas sclerotic WDLPS is more closely related to DDLPS. We propose that DDLPS arises from an MSP- like cell, sclerotic WDLPS from a C.APC-like cell, and adipocytic WDLPS from a pre-ADC-like cell (**Figure 3E**).

**Figure 3.**
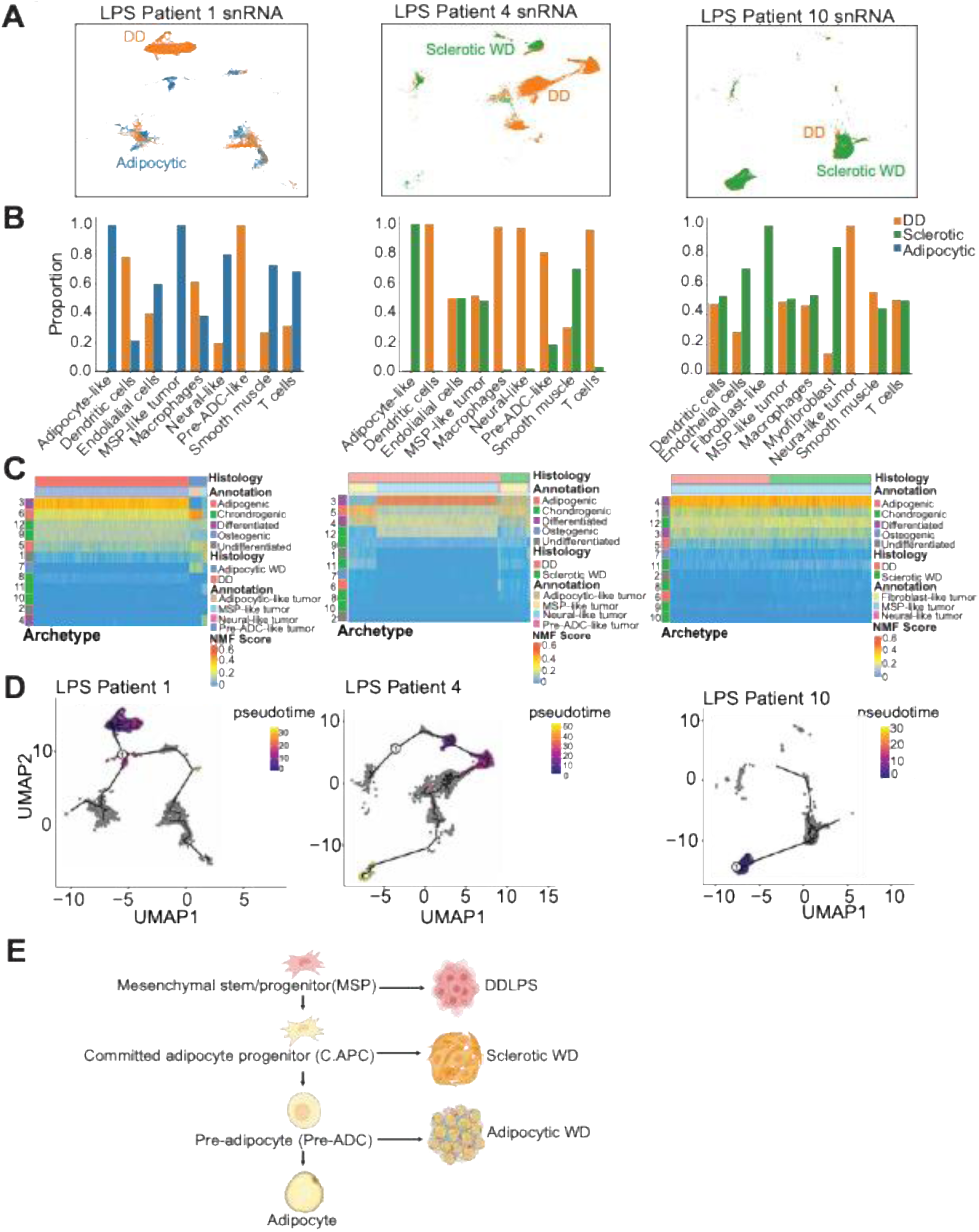
Matched WD-DD comparison reveals distinct drivers of WD and DD LPS components. (A) UMAP analysis of snRNA data showing annotations by pathologically annotated WD and DD LPS sample groups within three different tumors. (B) Proportion of cells by phenotype (from snRNA-seq data) within each tumor. (C) Lineage-specific archetype score heatmap comparing matched WD and DD samples. Row annotations indicate the lineages specified by each archetype. (D) Trajectory analysis of each individual tumor with the most differentiated cluster set as the root cells. (E) Schematic of findings from matched WD/DD analysis.

### Spatial transcriptomics-based tumor niches suggest subtype-specific tumor architecture

Given the intra-tumoral heterogeneity observed in paired WDLPS and DDLPS tumors, we next examined the spatial relationships using spatial transcriptomics. We applied the Xenium platform to capture single-cell-resolved spatial gene expression for LPS-relevant transcripts. Consistent with snRNA-seq results, spatial clustering identified adipocytic, progenitor-like, mesenchymal, and immune populations annotated using canonical markers (**Figure 4A-C, F; Supplementary Figure S8**). Cell type quantification revealed enrichment of adipocyte-like tumor clusters in WDLPS and progenitor-like clusters in DDLPS, indicating intrinsic differences in tumor cellular composition within matched samples (**Figure 4C**).

**Figure 4.**
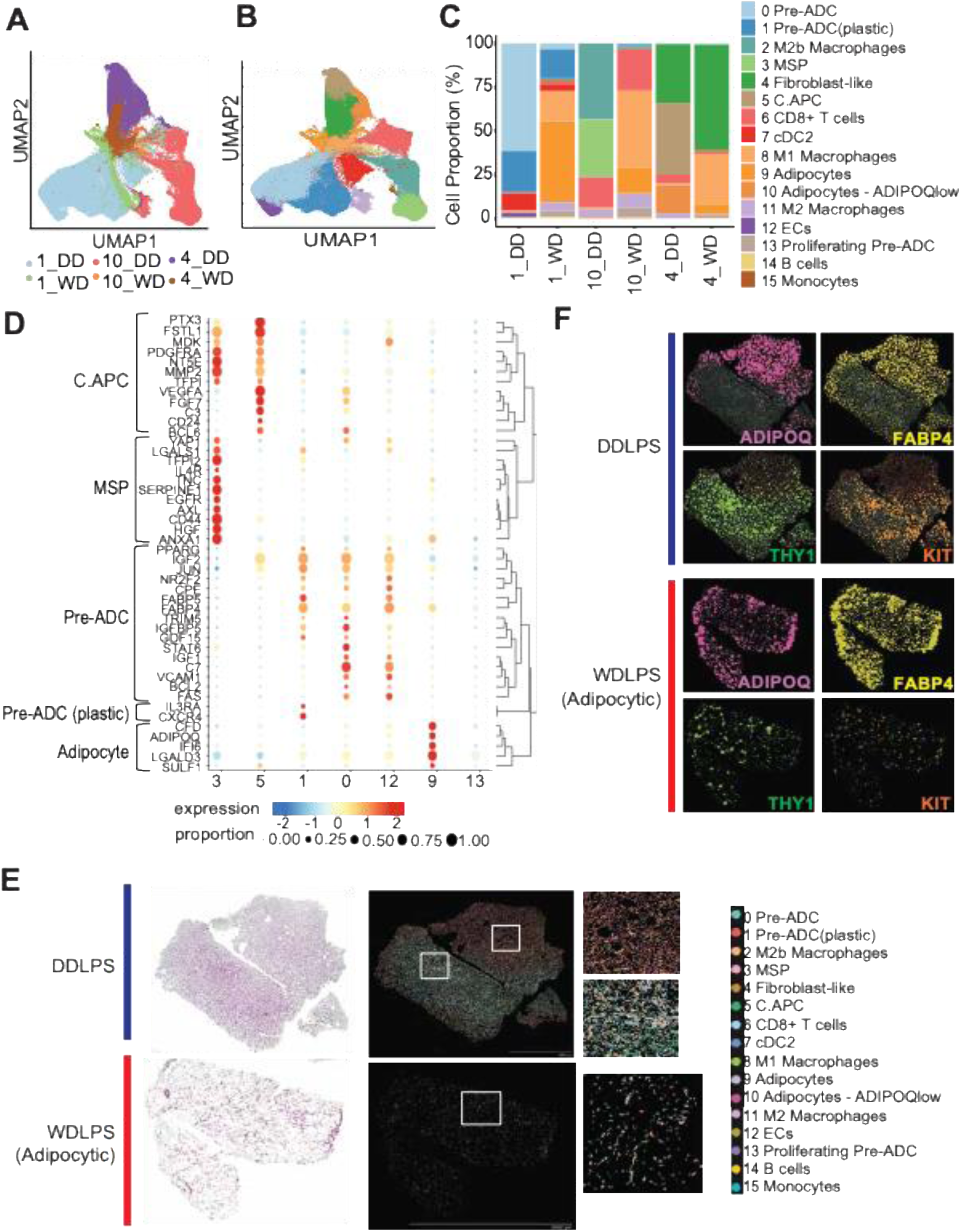
Spatial Mapping Reveals Subtype-Specific Tumor Architecture and Immune Engagement in LPS. (A) UMAP analysis of Xenium spatial transcriptomic data, projecting each of the 6 individual tumors as a different color. (B) UMAP analysis identified 16 clusters from Xenium spatial transcriptomic analysis of 6 WD/DD LPS matched pairs. (C) Stacked bar plot showing proportion of each cell type within each tumor. (D) Images of DDLPS and WDLPS tumor sections with H&E staining (left) and clusters identified (right). (E) Expression of adipocyte genes *ADIPOQ* and *FABP4* versus mesenchymal genes *THY1* and *KIT*, in DDLPS and matched adipocytic WDLPS. (F) Bubble plot showing defining genes for each tumor cluster.

Spatial projection of UMAP-derived clusters onto tissue sections confirmed that transcriptional heterogeneity corresponded to histological features and spatial organization. In one matched pair, the DDLPS component showed high predominance of MSP-like tumor cells with diffuse tumor–immune cell distribution (**Figure 4D-E**). In contrast, the matched WDLPS component contained distinct adipocyte-like tumor clusters, reduced mesenchymal markers (e.g. *THY1, KIT*), enriched adipocytic markers (e.g. *ADIPOQ, FABP4*), and more organized immune cell spatial patterns compared to the DDLPS component (**Figure 4D-E**).

Analysis of two additional matched W/DD LPS pairs revealed similar spatial patterns. In the second pair, the DDLPS component contained discrete tumor clusters expressing adipocyte/pre-ADC markers (*ADIPOQ, FABP4*) and early progenitor markers (*THY1, KIT*) (**Supplementary Figure 9A**). The matched WDLPS component retained these clusters but showed reduced spatial compartmentalization, minimal *THY1/KIT* expression and enrichment for adipocyte-like tumor cells expressing *ADIPOQ* and *FABP4* (**Supplementary Figure 9B**). In the third pair, the DDLPS region showed spatial exclusion of macrophages from *KIT*-high C.APC-like tumor clusters, while CD8⁺ T cells formed ring-like niches around fibroblast-like tumor cells that co-localized with M2 macrophages (**Supplementary Figure 10A**). In contrast, the WDLPS component contained adipocyte-like tumor cells and fibroblast-like cells, with lower-density CD8⁺ T cell niches, suggesting a more diffuse immune engagement (**Supplementary Figure S10B**). Together, these results demonstrate that WDLPS and DDLPS exhibit fundamentally distinct spatial transcriptional architectures, with adipocyte-like tumor populations predominating in WDLPS and progenitor- and mesenchymal-like populations dominating DDLPS, reflecting subtype-specific tumor architecture and cellular composition.

### Contrasting patterns of spatial niches harboring tumor immune microenvironment in LPS subtypes

To further characterize subtype-specific tumor–immune interactions in LPS, we compared immune compartments across WDLPS and DDLPS tumors and applied spatial niche analysis to resolve tumor–immune interactions in situ. Macrophages (69%) and T cells (23%) constituted the majority of immune cells and were most abundant in DDLPS, followed by sclerotic WDLPS and adipocytic WDLPS (**Figure 5A-C**). Eleven tumor-associated macrophage (TAM) clusters were identified and grouped by functional markers into interferon-primed (IFN; clusters 24, 35), resident tissue macrophage-like (RTM; clusters 8, 38), lipid-associated (LA; clusters 1, 39, 41), inflammatory cytokine-enriched (Inflam; clusters 0, 7), proliferating (Prolif; cluster 28), and regulatory (Reg; cluster 12) states (**Supplementary Figure S11**). DDLPS tumors contained higher macrophage abundance than WDLPS, with enrichment of IFN-, angiogenic-, and LA-associated TAM markers (**Figure 5B**). Sclerotic WDLPS differed from adipocytic WDLPS by increased RTM (especially HES1) and decreased angio TAM markers (especially VEGFA).

**Figure 5.**
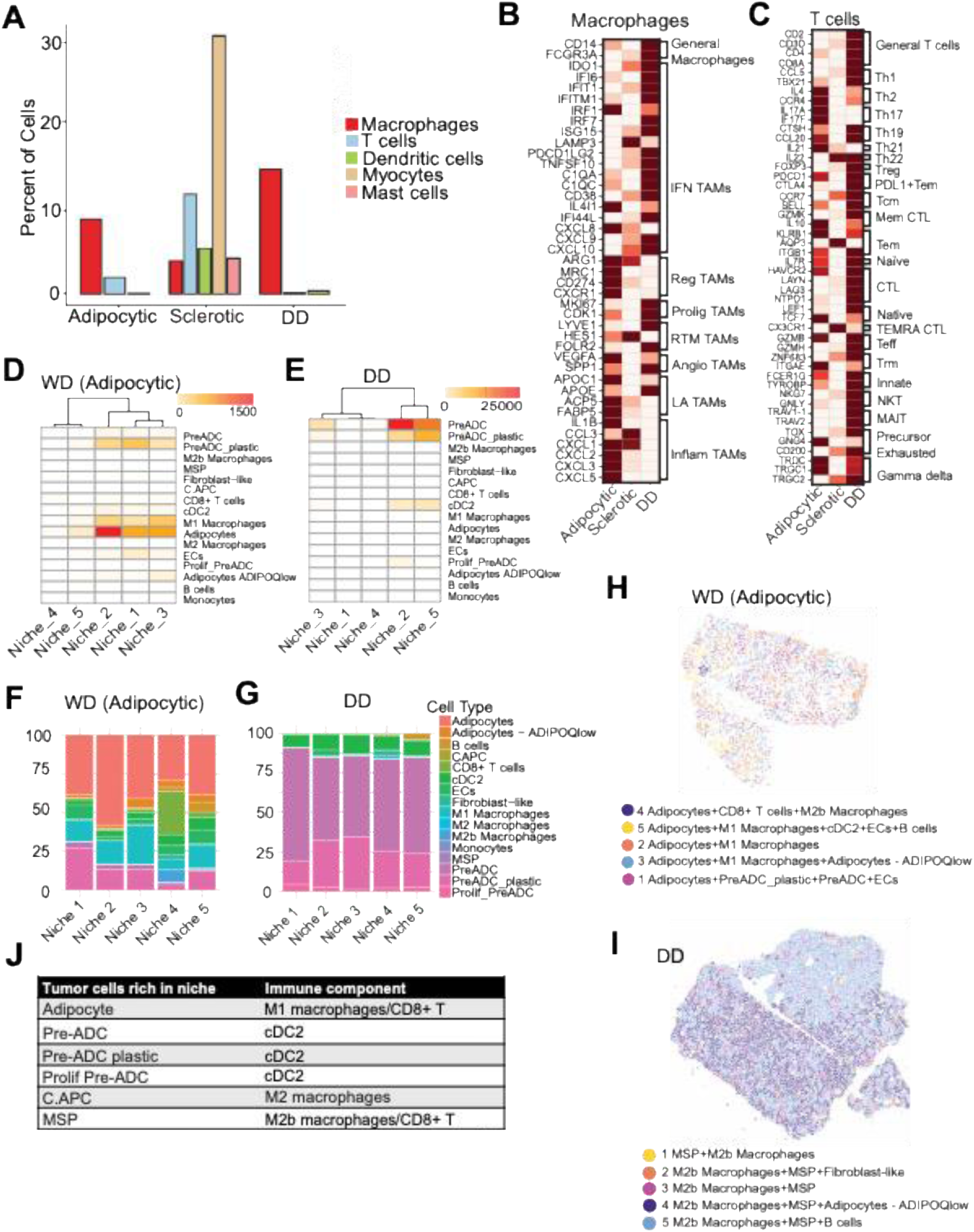
Distinct niches and microenvironments in WDLPS versus DDLPS. (A) Proportion of immune cells within adipocytic WDLPS, sclerotic WDLPS, and DDLPS tumors. (B) Heatmap showing markers of tumor-associated macrophage subtypes by adipocytic WDLPS, sclerotic WDLPS, and DDLPS. (C) Heatmap showing markers of tumor-associated T cell subtypes by adipocytic WDLPS, sclerotic WDLPS, and DDLPS. (D-E) Spatial maps of niches identified in WDLPS (D) and DDLPS (E) components from patient #1. (F-G) Stacked bar plots showing proportion of cells by cluster type that made up each niche within WDLPS (F) and DDLPS (G) components. (H-I) Niche composition of 5 niches identified in WDLPS (H) and DDLPS (I) components. (J) Chart summarizing the spatial co-localization of different tumor cell subsets (left column) with the most commonly co-localizing immune cell component (right column) from all three patients.

Five T cell clusters were identified, including two predominantly CD4⁺ and two predominantly CD8⁺ populations (**Supplementary Figure S11**). These were further classified as effector memory T cells (Tem, cluster 24, expression of *ITGB1*), naïve CD4+ (cluster 39, expression of *IL7R*) and cytotoxic T lymphocytes (CTL, cluster 38, expression of *HAVCR2/TIM3* and ENTPD1). The largest T cell cluster was a CD8+ cluster (cluster 4) that expressed markers of effector memory (*PDL1, CTLA4*), central memory (*CCR7, SELL*), and effector T cells (granzymes; **Supplementary Figure S11**). DDLPS tumors expressed higher levels of markers of exhausted T cells (*HAVCR2, LAYN, LAG3, ENTPD*1, *PDCD1, CTLA4*) compared to WDLPS (**Figure 5C**). Sclerotic WDLPS tumors exhibited higher Tem (*ITGB1*) and naïve (*IL7R*) signatures in comparison to adipocytic WDLPS and DDLPS. Adipocytic WDLPS tumors showed enrichment of Th2, Th17, and γδ T cells relative to both sclerotic WDLPS and DDLPS (**Figure 5C**).

We next performed spatial niche analysis on matched WDLPS–DDLPS tumors from three patients, revealing histology-specific tumor–immune architectures. In the first matched pair, DDLPS displayed structured niches with immune exclusion, whereas WDLPS exhibited immune-permissive states (**Figure 5D-I**). The WDLPS component was enriched for adipocyte-like tumor cells in niche 2, with plastic PreADC-like tumor cells distributed across tumor cell-rich niches (**Figure 5D, F, H**). Notably, dendritic cells were sparse, and M1 macrophages were enriched, consistent with a more pro-inflammatory environment. In contrast, the DDLPS component contained discrete niches dominated by pre-ADC tumor cells (Niche 1), alongside mixed niches containing both differentiated and plastic tumor states (Niches 3 and 5) (**Figure 5E, G, I**). cDC2-type dendritic cells were enriched in multiple DDLPS niches, indicating potential antigen presentation activity. Niche 5 contained PreADC cells and B cells, suggesting humoral immune involvement.

Similar patterns were observed in additional matched tumors. In patient 10, immune infiltration co-localized with mesenchymal and fibroblastic cell types in both WDLPS and DDLPS (**Supplementary Figure S9**). WDLPS niches contained adipocyte-like tumor cells co-localizing with M1 macrophages and CD8⁺ T cells (Niches 2,4, and 5), whereas a distinct niche (Niche 1) contained ADIPOQ-low adipocyte-like tumor cells with M1 macrophages but limited CD8⁺ T cell presence, indicating niche-specific immune modulation (**Supplementary Figure S9**). In the matched DDLPS component, MSP-like tumor cells dominated several niches (Niches 1, 3, and 5) that also contained M2b macrophages and CD8⁺ T cells, suggesting immune-suppressive tumor–immune engagement (**Supplementary Figure S9**). In the third matched tumor pair (patient 4), DDLPS exhibited immune-excluded C.APC niches surrounded by fibroblast-rich stromal regions containing T cells and M2 macrophages, consistent with stromal-mediated immune exclusion (**Supplementary Figure S10**). In contrast, WDLPS lacked C.APC-like populations and was dominated by adipocyte-like tumor cells, with M1 macrophages present across both adipocytic and fibroblast-rich regions, indicating a more inflammatory microenvironment (**Supplementary Figure S10**). Thus, spatial niche analysis across matched DDLPS and WDLPS tumors from three patients revealed distinct and previously unrecognized differences in the immune microenvironment, shaped by tumor cell differentiation state (**Figure 5J**).

### Epigenomic dissection of LPS reveals lineage-specific and dedifferentiation-associated regulatory programs

Next, we sought mechanistic insight into the observed underlying tumor cell plasticity, differentiation states, and lineage hierarchy in LPS. During normal homeostasis and tumorigenesis, each cellular state is characterized by unique epigenetic patterns and specific chromatin states, such as enhancers, are driver events.^30^ We performed snATAC-seq across tumor and stromal cell populations which enables discovery of open chromatin regions, including putative enhancers and associated regulatory TFs. Gene activity scores derived from accessibility peaks surrounding cell type–defining genes showed strong concordance with transcriptional signatures, indicating close correspondence between gene expression and epigenetic heterogeneity (**Figure 6A-B; Supplementary Figure S12A-B**). Unsupervised clustering identified accessibility-based cell populations that structured the UMAP manifold (**Supplementary Figure S12C**). Restricting analysis to malignant cells excluded immune-derived accessibility signatures and revealed finer stratification of tumor-intrinsic chromatin states across annotated cell subtypes and unsupervised clusters (**Supplementary Figure S12D–E**).

**Figure 6.**
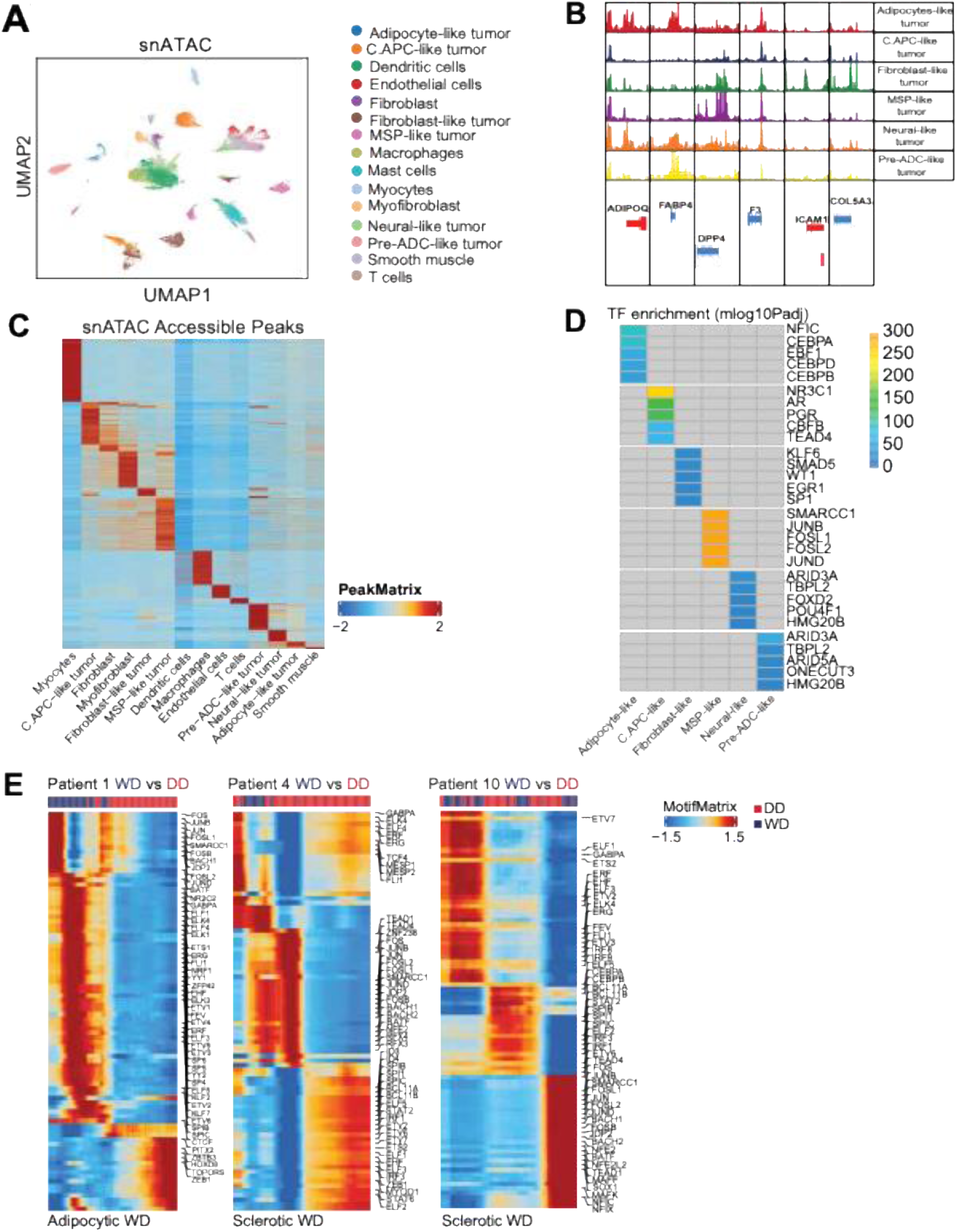
Regulatory circuitry governing LPS progression identified by integrative epigenomic and TF analysis. (A) UMAP analysis of snATAC-seq with annotations by cell type according to the snRNA-seq data. (B) snATAC-seq track plots for marker genes showing specific peak enrichments based on cell types and tumor de-differentiation stages. (C) Heatmap showing unsupervised clustering of snATAC peaks and differential peak enrichments based on cell types. (D) Heatmap showing unsupervised clustering of TF deviation scores and differential TF motifs enriched in the malignant cell clusters. (E) Pseudo-time trajectory analysis based on TF deviation scores showing shifts in regulatory TFs during progression from WDLPS to DDLPS.

Distinct cell type–specific chromatin accessibility patterns were observed, with characteristic peak enrichments distinguishing tumor and stromal compartments (**Figure 6C, Supplementary Figure S13**). TF motif enrichment analysis of cell type–specific accessible peaks further revealed divergent regulatory programs across tumor states (**Figure 6D; Supplementary Figure S14**). Early progenitor populations, including MSP cells, were enriched for TF motifs associated with developmental signaling and stress responses (SMARCC1, JUNB, FOSL1, FOSL2, JUND). C.APCs showed enrichment for NR3C1, AR, PGR, CBFB, and TEAD4 motifs, consistent with progenitor maintenance and nuclear hormone receptor signaling. Pre-ADC-like tumor clusters exhibited preferential accessibility at ARID3A motifs, implicating chromatin remodeling programs that may restrain terminal adipogenic differentiation (**Figure 6D**). In contrast, differentiated adipocyte-like clusters showed strong enrichment for CEBPA/B/D, classical regulators of terminal adipocyte identity and lipid metabolism. Fibroblast-like tumor cells showed enrichment for KLF6, SMAD5, WT1, EGR1, and SP1 motifs associated with mesenchymal differentiation and extracellular matrix remodeling. Neural-like tumor cluster exhibited enrichment for ARID3A, TBPL2, FOXD2, POU4F1, and HMG20B binding sites, suggestive of epigenetic-driven plasticity. Together, these findings highlight the hierarchical and dynamic nature of TF-mediated chromatin regulation across tumor cell states in LPS, supporting a model wherein tumor plasticity and lineage identity are epigenetically encoded.

We next inferred cell state transitions during LPS progression using gene regulatory network analysis of three matched WDLPS–DDLPS tumor pairs. Pseudotime ordering using scMEGA reconstructed trajectories from WDLPS to DDLPS and identified genome-wide regulatory dynamics associated with cell state transitions from WDLPS to DDLPS. Two trajectory patterns emerged. In patients 1 and 4, clear regulatory divergence was observed along the WDLPS–DDLPS continuum, with WDLPS enriched for AP-1 motifs and DDLPS enriched for mesenchymal regulators such as ZEB1 (**Figure 6E**). In contrast, patient 10 showed minimal regulatory divergence between WDLPS and DDLPS states.

To further characterize epigenetic regulation across LPS subtypes, we performed CUT&RUN profiling of key histone marks - H3K27Ac (active enhancer), H3K4me3 (active promoter), and H3K27me3 (repressive mark) - in one DDLPS and two WDLPS tumors (each with biological duplicates). Principal component analysis (PCA) of H3K27Ac peaks clearly separated WDLPS and DDLPS samples (**Figure 7A**). Replicates showed strong concordance, while DDLPS exhibited greater inter-sample divergence, suggesting increased regulatory variability (**Figure 7B**). Differential peak analysis identified 1,109 H3K27Ac peaks enriched in WDLPS and 676 peaks enriched in DDLPS (**Figure 7C**). Enhancer regions near adipogenic genes - including *CD34, F3, and ICAM1* - showed higher H3K27Ac signal in WDLPS, whereas DDLPS showed stronger H3K27Ac signal at loci associated with progenitor and mesenchymal regulators such as *KIT, PDGFRA*, and *YAP1* (**Figure 7D**).

**Figure 7.**
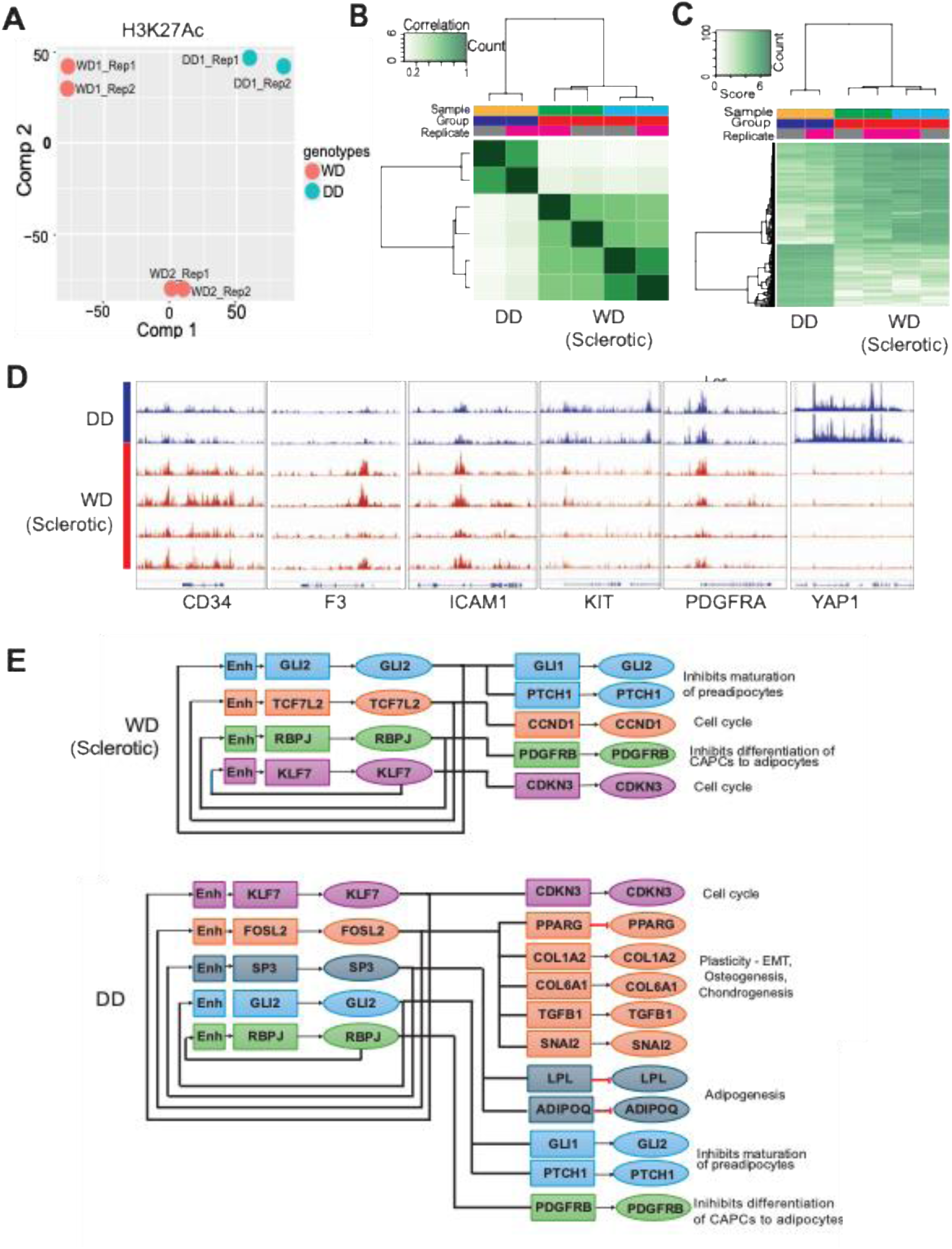
Differential enhancer activation in WD versus DD LPS. (A) PCA analysis of H3K27Ac CUT&RUN data. (B) Correlation heatmap analysis between biological replicates. (C) Differential peak analysis identified 1109 H3K27Ac peaks enriched in WDLPS and 676 peaks enriched in DDLPS. (D) Track plots of CUT&RUN analysis showing H3K27Ac enrichment at CD34, F3, ICAM1, KIT, PDGFRA, YAP1 loci comparing the bigwig tracks of DDLPS and WDLPS samples (n=3) (E) Schematic showing the core regulatory circuits identified in sclerotic WDLPS and DDLPS with corresponding target genes.

Similar subtype separation was observed for promoter-associated H3K4me3 profiles (**Supplementary Figure S15A**). Replicate correlations confirmed high reproducibility and greater promoter variability in DDLPS (**Supplementary Figure S15B**). Differential binding analysis revealed 3,630 promoter peaks gained in WDLPS and 291 in DDLPS (**Supplementary Figure S15C**). Promoters of key adipogenic regulators, including *PPARG*, showed increased H3K4me3 in WDLPS, whereas progenitor-associated genes, such as *KIT*, were selectively marked in DDLPS (**Supplementary Figure S15D**). H3K27me3 (repressive mark) profiles also distinguished WDLPS from DDLPS, with subtype-specific clustering (**Supplementary Figure S14E**). While DDLPS replicates were tightly correlated, WDLPS samples displayed greater epigenetic variability (**Supplementary Figure S14F**). Differential peak analysis identified 131 promoter regions with increased H3K27me3 in WDLPS, and only 18 enriched in DDLPS, suggesting differential silencing mechanisms across subtypes (**Supplementary Figure S14G**). For example, OSBPL8, an adipogenic gene, showed H3K27me3-mediated silencing in DDLPS, while CD6 was repressed in WDLPS (**Supplementary Figure S14H**). Together, these profiles corroborate the chromatin accessibility landscape and reveal subtype-specific enhancer activation, promoter regulation, and transcriptional repression.

During normal development, a small group of core TFs defines cell identity by regulating themselves and one another through stabilizing feedback loops, thus forming the core transcriptional regulatory circuitry (CRC).^31^ To identify CRCs in LPS, we integrated snATAC-seq, snRNA-seq, and CUT&RUN H3K27ac profiling (**Supplementary Figure S16A**). We first identified the top TF motifs enriched in open chromatin regions from tumor-derived snATAC clusters across adipocytic and sclerotic WDLPS and DDLPS samples followed by comparison of their expression (top 50 TFs ranked by motif enrichment) to assess group-specific differences. TFs showing both strong motif enrichment and upregulated expression in sclerotic WDLPS or DDLPS were further examined for enhancer gains and putative self-regulatory binding sites using CUT&RUN data. TFs meeting all criteria were retained, and we next evaluated the expression of their predicted direct targets (**Supplementary Figure S16B**). In adipocytic WDLPS, several adipogenic TFs were upregulated; however, self-regulatory circuitry analysis could not be completed due to the absence of CUT&RUN data for these technically challenging samples. In sclerotic WDLPS, we identified three CRC-candidate TFs (GLI2, TCF7L2, and RBPJ) and in DDLPS three TFs (KLF7, FOSL2, and SP3), that ranked among the top 50 enriched motifs and exhibited self-regulatory enhancer occupancy (**Supplementary Figure S17A**). Notably, KLF7 was also highly expressed in sclerotic WDLPS, whereas RBPJ and GLI2 were highly expressed in DDLPS, suggesting partially overlapping regulatory programs between these states.

Analysis of direct target gene expression further highlighted upregulation of cell-cycle regulators (CCND1, CDKN3) and inhibitors of adipocyte differentiation (GLI1, PTCH1, PDGFRB) in sclerotic WDLPS. In DDLPS, we additionally observed increased expression of TF targets associated with epithelial–mesenchymal transition (SNAI2) and osteo/chondrogenic programs (COL1A2, COL6A1, TGFB1), consistent with mesenchymal progenitor plasticity, along with downregulation of adipogenic maturation genes (PPARG, LPL, ADIPOQ) (**Figure 7E; Supplementary Figure S17B**).

Collectively, these results indicate that sclerotic WDLPS and DDLPS are driven by distinct but partially overlapping CRC programs that promote proliferation, suppress adipogenic maturation, and sustain mesenchymal lineage plasticity, providing a mechanistic framework for dedifferentiation and aggressive tumor behavior in LPS.

## DISCUSSION

Together, these findings demonstrate that WDLPS and DDLPS tumor cells lie along a continuum of adipocytic differentiation, with WDLPS retaining more adipocyte-like features and DDLPS displaying transcriptional signatures consistent with early mesenchymal progenitor or undifferentiated states. Comparison of adipocytic and sclerotic WDLPS subtypes further revealed that sclerotic tumors display greater lineage plasticity and transcriptional profiles more closely aligned with DDLPS, including enrichment of chondrogenic and osteogenic archetypes and increased expression of progenitor markers. Despite their histological classification as WDLPS, these tumors likely represent an intermediate state with increased propensity for progression toward dedifferentiation. These observations suggest that histological classification alone may underestimate the molecular heterogeneity and plasticity within WDLPS tumors. Prior work has ascribed a degree of developmental maturation to LPS by comparing gene expression profiles of LPS tumors to different timepoints of MSCs differentiating into adipocytes.^32^ It will be of future interest and importance to determine whether lineage state better predicts clinical outcomes than the current binary classification of WDLPS versus DDLPS.

Our spatial analyses further reveal that tumor cell differentiation state shapes immune–tumor interactions within the LPS microenvironment. In DDLPS, immune activity was largely confined to peripheral or mixed niches, whereas C.APC-dominated regions exhibited immune exclusion. In contrast, WDLPS tumors showed broader immune infiltration across niches containing adipocyte-like or plastic tumor cells, with pro-inflammatory M1 macrophages as prominent components. Distinct tumor cell states preferentially recruited specific immune populations: adipocyte-rich niches were associated with M1 macrophages and CD8⁺ T cells, whereas pre-ADC–rich niches preferentially recruited cDC2 dendritic cells. C.APC-like tumor regions were enriched for M2 macrophages, and MSP-like niches showed co-enrichment of M2b macrophages and CD8⁺ T cells. These interactions were accompanied by cell state–specific inflammatory programs. Adipocyte-rich niches expressed *LGALS3* and *IFI6*, whereas pre-ADC niches co-expressed pro-inflammatory mediators (*VCAM1, IGF1, JUN*) alongside regulatory factors (*PPARG, BCL6*). C.APC-like niches displayed angiogenic and wound-healing signals (*VEGFA, PDGFRA, MDK*) together with immunosuppressive mediators (*FSTL1, NT5E*). MSP-like niches expressed invasive and immunomodulatory genes (*AXL, YAP1, MMP2, IL4R, LGALS1*). Together, these results highlight spatially organized functional heterogeneity within the LPS microenvironment and support a model in which tumor cell state shapes immune architecture.

Recent single-cell studies have similarly advanced understanding of tumor plasticity and lineage relationships in WD/DD LPS.^23,24^ One analysis identified adipocyte stem-cell–like progenitors carrying ancestral genomic alterations shared by WDLPS and DDLPS components, suggesting early divergence of the two lineages rather than linear progression from WDLPS to DDLPS.^24^ In that model, DDLPS retains stem-like adipocyte progenitor features, while adipogenic differentiation is suppressed by a TGF-β–rich immunosuppressive microenvironment. Complementary work integrating snMultiome and spatial profiling identified loss of IGF1 signaling and downregulation of the adipocyte-specific *PPARG2* axis as key features of dedifferentiation.^33^ Further, DDLPS displays reactivation of early mesenchymal/development programs and heightened sensitivity to IGF1R inhibition. Together, our work along with these studies support a model in which DDLPS arises from adipocytic progenitor-like cells that diverge early into WDLPS and DDLPS trajectories and are maintained by epigenetic and metabolic reprogramming (via IGF1/PPARG2 axis) rather than simple dedifferentiation of mature adipocytic cells. These findings suggest potential therapeutic strategies targeting TGF-β signaling or IGF1R pathways in DDLPS.

The integration of snRNA-seq and snATAC-seq enabled simultaneous profiling of transcriptional and epigenomic states within individual tumor cells, allowing direct linkage of chromatin accessibility with transcriptional programs. This approach revealed regulatory networks and transcription factor (TF) motifs associated with WDLPS and DDLPS states and transitions between them. - a question that has heretofore not been adequately addressed. Integration with chromatin state profiling further suggests that rewiring of core transcriptional regulatory circuitry (CRC) represents a central mechanism driving the transition from well-differentiated to dedifferentiated states. The discovery of GLI2, TCF7L2, and RBPJ in sclerotic WDLPS and KLF7, FOSL2, and SP3 in DDLPS highlights subtype-specific regulatory dependencies. The overlap between sclerotic WDLPS and DDLPS TF circuits suggests that dedifferentiation may arise not solely from the accumulation of genetic changes but also from the reinforcement of shared epigenetic and transcriptional programs that sustain proliferation and permit phenotypic plasticity. The involvement of TFs linked to Hedgehog, Notch, AP-1, and KLF pathways highlights actionable regulatory nodes with existing pharmacologic modulators. These findings suggest that targeting enhancer-driven TF circuits or restoring adipogenic differentiation may therefore represent promising therapeutic strategies that warrant functional validation in patient-derived models.

This study has several limitations. First, the cohort size was relatively small (23 samples from 13 patients), and larger studies will be required to determine the generalizability of these findings. Second, trajectory analyses assumed that DDLPS evolves from a WDLPS precursor, although these states may arise independently from shared progenitor populations. Finally, while we define the differences in lineage plasticity, tumor-immune interactions and core regulatory epigenetic programs associated with WDLPS (adipocytic and sclerotic) and DDLPS, the definitive proof for predicted association requires functional genetic experiments in relevant immune proficient models (that do not exist). Hence, these studies will be the focus of our future studies.

In summary, we present a spatially resolved single-nucleus multiome atlas of primary untreated WD/DD LPS that integrates transcriptional and epigenomic profiling at single-cell resolution. These data reveal lineage hierarchies, spatial tumor–immune interactions, and regulatory circuits that underlie tumor plasticity and dedifferentiation. Our findings identify candidate regulatory dependencies and therapeutic targets that warrant further investigation in WD/DD LPS, a disease with substantial unmet clinical need.

## METHODS

### Patients and ethics

This study was approved by the University of Texas MD Anderson Cancer Center Institutional Review Board (protocols PA12-0305, 2022-0278, and LAB04-0890) and was conducted in accordance with the U.S. Common Rule and the Declaration of Helsinki. Clinical and genomic data were obtained following signed informed consent onto prospective institutional protocols or under retrospective review protocols with a limited waiver of authorization, as this is a retrospective project that involves no diagnostic or therapeutic intervention and no direct patient contact. Clinical characteristics, including demographic characteristics, disease-associated characteristics (e.g. tumor size, site, etc), and survival were collected from patient electronic health records.

### Patient specimens

Specimens from patients who underwent surgical resection for WD/DD LPS were utilized. The patients had not received chemotherapy or radiation prior to surgery, so the specimens used in this study were treatment-naïve. An expert sarcoma pathologist reviewed adjacent H&E-stained slides to confirm the identity as either WDLPS or DDLPS. WDLPS was further classified as “sclerotic” or “adipocytic,” as previously described.^16^ In patients, sclerotic tumors were defined as > 10% sclerosis, and adipocytic tumors were defined as < 10% sclerosis. The particular tissue samples used in this analysis were composed predominantly (>90%) of sclerotic or adipocytic cells.

### Nuclei isolation for single-cell sequencing

Nuclei isolation was performed according to the 10x Genomics “Demonstrated Protocol Nuclei Isolation from Embryonic Mouse Brain for Single Cell Multiome ATAC + Gene Expression Sequencing,” CG000366 RevC. Briefly, tissue specimens were thawed on ice and minced with a scalpel or razor blade. The minced tissue was added to a 1.5 mL microcentrifuge tube with 500 μL 0.1X lysis buffer, homogenized using a pellet pestle, and incubated on ice for 10 min. Cell/tissue suspensions were mixed 10x with a wide-bore pipette and transferred to a 100 μm Flowmi Cell Strainer and washed with 500 μL wash buffer. Cell suspensions were centrifuged (500g for 5 min at 4 °C) followed by gentle removal of the supernatant without disrupting the cell pellet, another wash with 1 mL wash buffer, centrifuged, and resuspended in wash buffer, and passed through a 40 μm Flowmi Cell Strainer. The concentration and quality of nuclei were assessed using 0.4% trypan blue staining and verification on a hemocytometer. The nuclei were centrifuged again and resuspended in an appropriate volume of chilled 1X Nuclei Buffer (10x Genomics, 2000153) to a concentration of 8000 nuclei/μL. Nuclei were centrifuged again, resuspended in chilled 1X Nuclei Buffer (10x Genomics, 2000153) to 8000 nuclei/μL, and immediately used to prepare snATAC-seq and snRNA-seq libraries.

### Library preparation for single nucleus multiome

Single nucleus multiome sequencing was performed using 10x Genomics Chromium Next GEM Single Cell Multiome ATAC + Gene Expression Reagent Bundle (10x Genomics) following the manufacturer’s protocol “CG000338 Rev E Chromium Next GEM Single Cell Multiome ATAC + Gene Expression.” The main steps of this protocol include: (1) nuclei transposition, (2) GEM generation and barcoding, (3) post-GEM incubation cleanup, (4) pre-amplification PCR, (5) ATAC library construction, (6) cDNA amplification, and (7) gene expression library construction. Briefly, nuclei suspensions from previous step with targeted recovery of 10,000 nuclei per sample were mixed and incubated (37 °C for 60 min) with transposition mix. Also, during this step, adapter sequences are added to the end of the DNA fragments. Next, the transposed nuclei were mixed with a master mix containing barcoding reagents and loaded onto a Chromium Next GEM Chip J along with Single Cell Multiome GEX Gel Beads and partitioning oil. Gel beads-in-emulsion (GEMs) were generated by delivering nuclei at a limiting dilution to the gel beads using the 10x Chromium Controller. The GEMs were captured, cleaned up, then dissolved, releasing oligonucleotides containing an Illumina P5 sequence, a 16-nucleotide 10x barcode for ATAC, and a spacer sequence. Primers containing an Illumina TruSeq Read 1 primer, a 16-nucleotide 10x barcode for GEX, a 12-nucleotide unique molecular identifier (UMI), and a 30-nucleotide poly(dT) sequence were also released during this step. These oligonucleotides were mixed with the nuclear lysate containing transposed DNA fragments, mRNA, and a master mix containing reverse transcriptase, generating the 10x barcoded transposed DNA for snATAC-seq and 10x barcoded cDNA for snRNA-seq. The reaction was quenched with a quenching reagent. GEMs were then broken, and pooled fractions were recovered using Silane magnetic beads to purify barcoded products. Barcoded transposed DNA and cDNA were amplified to fill in gaps for subsequent ATAC library construction and cDNA amplification. Next, the snATAC-seq library was constructed by adding a P7 sample index and performing PCR. Attention was turned to creating the snRNA-seq library, first by amplifying cDNA by PCR to generate sufficient material for library construction, then fragmented and size selected using SPRIselect (Beckman Coulter) to optimize the cDNA amplicon size. P5, P7, i7, and i5 sample indexes, and TruSeq Read 2 primer sequences are added via end repair, A-tailing, adaptor ligation, and PCR, followed by purification with SPRIselect and elution of final libraries.

Next, the libraries were checked for the fragment size distribution using Agilent 4200 Tape Station HS D1000 Assay (Agilent Technologies) and quantified with Qubit Fluorometric dsDNA Quantification kit (Thermo Fisher). The libraries were sequenced at the MD Anderson Advanced Technology Genomics Core facility, with each library on a separate lane of HiSeq4000 flow cell (Illumina), with the sequencing targeting above the minimum of 25,000 read pairs per nucleus sequencing depth, format of 100nt and parameters (Read 1—50 cycles, Read 2—50 cycles; with exception while sequencing together with snRNA libraries Read 1—100 cycles, Read 2—100 cycles).

### Analysis of snRNA-seq and snATAC-seq data

Raw Single Cell Multiome ATAC + Gene Expression data were pre-processed using Cell Ranger ARC v2.0.0 from 10x Genomics. This process included demultiplexing cellular barcodes, aligning reads, and generating both the gene count matrix and the aligned fragments file. For read alignment, the human reference genome GRCh38 (hg38) was used. Comprehensive quality control metrics were generated and assessed, with cells being meticulously filtered to ensure high-quality data for downstream analyses. In summary, for basic quality filtering, cells with low-complexity libraries (where detected transcripts were aligned to fewer than 200 genes, such as cell debris, empty drops, and low-quality cells) were excluded from further analysis. Additionally, cells likely in the process of dying or undergoing apoptosis (with more than 15% of transcripts derived from the mitochondrial genome) were also removed. Cells with high-complexity libraries (where detected transcripts were aligned to more than 15,000 genes) were excluded. Doublets or multiplets were identified using the DoubletFinder algorithm.^34^ The expected proportion of doublets was estimated based on cell counts and 10x Genomics expected multiplet rates. Data normalization using scTransform was then performed on the filtered gene–cell matrix with Seurat.^35^ Because single-cell sequencing technologies often generate numerous zero values or dropouts, we employed SAVER^36^ to recover gene expression from noisy and sparse snRNA-Seq data. The FindVariableFeatures function of Seurat was used to identify highly variable genes for unsupervised cell clustering. Principal Component Analysis (PCA) was conducted on the top 2,000 highly variable genes. An elbow plot, created with Seurat’s ElbowPlot function, was used to determine the number of significant principal components. The FindNeighbors function in Seurat was utilized to build the shared nearest neighbor graph based on the unsupervised clustering results obtained with the FindClusters function. Several iterations of clustering and sub-clustering analyses were conducted to identify major cell types and distinct transcriptional states. Dimensionality reduction and 2D visualization of the cell clusters were carried out using uniform manifold approximation and projection (UMAP)^37^ through the RunUMAP function, with the number of principal components for embedding matching those used in clustering. Differentially expressed genes for the clusters were identified using the FindAllMarkers function, with criteria of an FDR-adjusted P value < 0.05 and a log2(fold change) > 1.2.

We converted the Seurat object to a Scanpy-compatible format for downstream snRNA-seq analysis. All subsequent analyses were performed using Scanpy, and UMAP visualizations for different annotations were generated in Python 3.13. Different gene markers for each cell type and cluster were used to construct dot plots comparing the gene expression levels and gene activity scores.

Large-scale copy number variation (CNV) was inferred from snRNA seq data using inferCNV. We constructed a genes × cells count matrix and provided a gene position file (gene symbol, chromosome, start, end; hg38) sorted by genomic order. Endothelial cells were used as a normal reference. We performed inferCNV on all chromosomes except for chrX, chrY, and chrM. For UMI data, we set cutoff=0.1, enabled group-wise clustering (cluster_by_groups=TRUE), denoising (denoise=TRUE), and HMM segmentation (HMM=TRUE). The pipeline outputs cell-level CNV profiles and discrete gain/loss calls; we summarized per-cell CNV burden (variance of the smoothed CNV signal/fraction of altered bins) and used coherent arm-level events to demarcate malignant versus non-malignant populations. Chromosome 12 inferCNV scores were used to identify chromosome 12q amplification, which is present in WD/DD LPS cells.

Cellular differentiation state was quantified with CytoTRACE^38^ using raw UMI. CytoTRACE computes a gene-diversity–weighted signature that assigns a continuous score per cell (higher values indicate less differentiated, stem-like states). Scores were integrated into the single-cell object for visualization on UMAP and compared across clusters. Associations between CytoTRACE and CNV burden were evaluated by Spearman correlation, and enrichment of known stemness and lineage markers was assessed for biological validation.

The R package ArchR^28^ was used for snATAC-seq data processing and analysis. Cells with a transcription start site (TSS) enrichment score of less than 45 and fewer than 1,000 unique fragments were filtered out. Doublets were inferred and removed using standard ArchR parameters. Dimensionality reduction was conducted using Iterative Latent Semantic Indexing,^29^ and single-cell embeddings were generated using UMAP. Peaks were identified using the MACS2 algorithm.^30^ Further downstream analysis, including cell label predictions, gene score calculation, marker peak identification, and motif enrichment analysis with ChromVAR was annotated by using Cis-BP motif sets, using ArchR’s default analytic functions. We utilized cross-modality integration methods, specifically the FindTransferAnchors function within ArchR, to align and integrate data from different modalities. This process was further refined by correcting the integration using shared 16-nucleotide barcode identifiers, ensuring accurate alignment and correlation between the datasets.

The Snapatac2 python package was used for processing and analyzing snATAC seq data. Cells with a TSS enrichment score below 5 and fewer than 1,000 unique fragments were excluded to ensure high-quality nuclei. Potential doublets were detected and removed using Scrublet, which identifies artificial multiplets by simulating random pairings of chromatin accessibility profiles. Dimensionality reduction was performed using spectral embedding to obtain a lower-dimensional representation of chromatin accessibility patterns across single cells. Clustering was then conducted using a graph-based k nearest-neighbor approach, allowing identification of distinct cell populations based on accessibility similarity. Peaks in the matrix were identified by using the MACS3 algorithm. A gene activity matrix was generated by aggregating accessibility profiles across cells within each cluster and calculating z-scores to determine significantly enriched peaks. Finally, motif enrichment analysis was carried out using the Cis-BP transcription factor (TF) motif database to annotate regulatory elements associated with each cluster.

### Pseudotime trajectory inference with Monocle3

We performed pseudo-time trajectory analysis in pairwise (i.e. matched WDLPS and DDLPS) for patients 1, 4, and 10 to characterize dynamic transcriptional changes. A Monocle3 cell_data_set object was constructed from the normalized gene expression matrix with corresponding cell and gene metadata. Dimensionality reduction was performed using PCA followed by UMAP, after which a principal graph was learned on the low-dimensional embedding using learn_graph. Cells were ordered along the inferred trajectory using order_cells, with the root state manually defined by chosen cell types. We chose pre-ADC-like tumor cells for patient 1 and 4 and MSP-like tumor cells as root cells for patient 10. Pseudotime values were then used to visualize patient-specific tumor trajectories, with tumor cells colored by pseudotime and non-tumor cells shown in gray for reference when applicable.

### Gene regulatory network inference

Gene regulatory network inference was performed using the scMEGA R package, which integrates single nucleus multiome (snMultiome) data to link TF activity with gene expression dynamics. To begin, snRNA-seq and snATAC-seq datasets were integrated by co-embedding the two modalities into a shared low-dimensional space using canonical correlation analysis implemented in Seurat. A differentiation trajectory was then reconstructed to capture the transcriptional continuum from WD to DD LPS states. To identify key regulatory factors driving this progression, TF activity scores were computed for each cell based on motif accessibility. Motif matching was performed on accessible chromatin regions, and chromVAR was used to calculate deviation scores that quantify variability in motif accessibility across cells. These scores, coupled with gene expression profiles, were used by scMEGA to infer TF–target gene regulatory relationships along the LPS differentiation trajectory. To further characterize subtype-specific regulatory programs, we analyzed patients 1, 4, and 10 separately to distinguish between WDLPS and DDLPS. For each patient, we visualized the enriched TF motifs along the inferred differentiation trajectory by plotting heatmaps of motif activity scores, highlighting dynamic changes in regulatory element accessibility during tumor progression.

### Gene expression archetype analysis

We applied a Normalized Non-negative Matrix Factorization (N-NMF) algorithm to define transcriptional programs or “archetypes,” which capture the variability of gene expression patterns across the dataset in a low-dimensional subspace, as previously described.^29^ This approach is a semi-supervised machine learning approach, where we first trained N-NMF archetype coefficients on the MSC differentiation dataset and later used these trained archetypes to score the LPS datasets. N-NMF archetype coefficients were computed using an iterative algorithm to compute a gene-archetype coefficient matrix and cell-archetype score matrix (normalized to sum to 1 for each cell). Scoring of the LPS dataset with trained archetypes was then computed using a similar iterative algorithm, but with the gene-archetype coefficient matrix held fixed as determined in the MSC differentiation dataset. Heatmaps and composition bar plots were produced using the R packages ComplexHeatmap and ggplot2.

### CUT & RUN-seq and analysis

Nuclei from WDLPS and DDLPS specimens with single nuclear multiome data were isolated using the 10X genomics protocol (as above) except resuspended in nuclei cryopreservation buffer (10 mM Tris-HCl pH 7.4, 150 mM NaCl, 1% BSA, 1 mM DTT, 1 U/μl RNase inhibitor, 0.1% Tween 20, 20% glycerol) at 5-10 million nuclei/ml and slow frozen (-1°C/min) to -80°C. CUTANA™ CUT&RUN (*EpiCypher* #14-1048) was performed using an automated procedure (autoCUT&RUN) based on previous protocols.^42^ All steps were performed on Tecan Freedom EVO robotic platforms at ambient temperature unless indicated, with gentle shaking during incubations and magnetic capture for buffer exchange/washing. Appropriate reactions contained the SNAP-CUTANA™ K-MetStat spike-in panel to monitor antibody performance (*EpiCypher* 19-1002).

In brief, CUT&RUN^43^ was performed with nuclei (6,000 – 50,000 per reaction) immobilized on Concanavalin A magnetic beads (Con-A; *EpiCypher* #21-1401), 0.5 μg antibody (**Supplementary Table S5**), and overnight incubation at 4°C. After the addition and activation of pAG-MNase (*EpiCypher* 15-1016), CUT&RUN-enriched DNA was purified as per kit instructions or using SPRIselect beads (*EpiCypher* 21-1403) and quantified by PicoGreen (*ThermoFisher* P7589). Sequencing libraries were prepared with 5 ng DNA (or total recovered if less) and the CUTANA CUT&RUN Library Prep kit (*EpiCypher* 14-1001). Libraries were analyzed by Agilent TapeStation, pooled to equivalence, and sequenced (Illumina NexSeq 2000), targeting 7-10 million paired-end reads (PE100) per reaction.

Data analysis was performed on the high-performance computing cluster at MD Anderson (http://hpcweb.mdanderson.edu/citing.html). FastQC and MultiQC was used to assess quality of raw fastq reads, followed by alignment to hg38 reference genome by Bowtie2 using the following parameters: --end-to-end --very-sensitive --no-mixed --no-discordant -I 10 -X 700 –dovetail and adaptors were trimmed. Duplicate reads were removed using SAMBLASTER and converted to BAM files. Uniquely mapped reads for each target were normalized by total reads, down-sampled to 8 million reads per sample, sorted and indexed using Samtools^44^ version 0.1.19. Peaks were called using MACS2 and bigwig files generated using deepTools^45^ version 2.4.0 and viewed in Integrative Genomics Viewer.^46^

### Xenium In Situ spatial transcriptomics and data analysis

The 10x Genomics Xenium platform was utilized for spatial transcriptomics. We designed a custom 480-gene panel using differentially expressed genes from the snRNA-seq data and well-established markers of different immune cell subsets. A list of genes is shown in **Supplementary Table S6**. Using tissue from the three available matched WD/DD pairs, 5 mm formalin-fixed, paraffin-embedded sections were placed onto a Xenium slide. Tissues underwent deparaffinization and permeabilization, then the custom probe panel at 10 nM concentration was hybridized overnight at 50 °C. Slides were washed, then probes were ligated to seal the junction between probe regions that had hybridized to RNA at 37 °C for 2 hours, followed by amplification of ligation products at 37 °C for 2 hours. Tissues were washed, chemically quenched of background fluorescence, and nuclei stained with DAPI. Slides were loaded onto a Xenium Analyzer. The Xenium cell segmentation algorithm used DAPI images that were acquired and calculated in a 15-μm radius from the nucleus outward or until another cell boundary was reached. The on-instrument pipeline output included files containing the feature-cell matrix, the transcripts, and a CSV file of the cell boundaries (a differentially enriched gene list for each cluster). Further downstream analysis and data integration for all samples were performed off-instrument and visualized using 10x Genomics Xenium Explorer (version 3). After completing the run, H&E staining was performed on the slide. A Keyence microscope was used to obtain images of the stained slide.

Data were processed using the Seurat package in R and filtered to exclude cells with zero molecule counts. Normalization was performed with SCTransform to correct for technical variability, and all samples were merged into a single Seurat object for integrated analysis. After this, the combined dataset was normalized, scaled, and subjected to principal component analysis (PCA) followed by UMAP for dimensionality reduction using the top 30 components. Clustering was performed with a shared nearest neighbor algorithm at a resolution of 0.3 to identify transcriptionally distinct groups. Prior to marker detection, the PrepSCTFindMarkers was applied to ensure compatibility of SCT-transformed data with downstream differential expression analysis. Cluster-specific marker genes were identified using FindAllMarkers with a minimum expression threshold of 25% and a log2 fold change cutoff of 0.25. The top 100 marker genes per cluster were exported for downstream analysis.

Spatial niche analysis was performed on the Xenium spatial transcriptomics dataset using Seurat. Cell identities were first defined based on unsupervised clustering, with cluster assignments stored in the Seurat clusters metadata field. All cells present in the selected field of view were included in downstream niche analysis. A niche assay was constructed using the BuildNicheAssay function. For each cell, a local neighborhood was defined by identifying its 30 nearest spatial neighbors (neighbors.k = 30). Neighborhoods were characterized by the composition of Seurat clusters within each local neighborhood and subsequently grouped into 15 distinct spatial niches (niches.k = 15) based on similarity in cluster composition. Spatial distributions of both cell types and inferred niches were visualized using ImageDimPlot, to projects cells on to their original tissue coordinates. Cells were colored either by their Seurat cluster identity or by their assigned niche identity. Custom color palettes were applied to enhance visual distinction between niches. Further, to ensure the data integrity and absence of any missing values, a contingency table was generated between cell-type clusters and spatial niches to summarize the number of cells from each Seurat cluster within each niche. This table was converted to a matrix format and exported as a CSV file for downstream statistical analysis. Finally, the processed Seurat object containing niche assignments with metadata formation was saved as an RDS file for downstream analyses.

### Code and data availability

Relevant code used for data processing and analysis of N-NMF are available at https://github.com/Ludwig-Laboratory/sarcoma_differentiation_landscape. Code used for multiome, CUT&RUN, and spatial transcriptomics are available at https://gitlab.com/railab/liposarcoma. Datasets used int his study are available in the Gene Expression Omnibus with the following identifiers: MSC differentiation (GSE241130), single cell multiome (GSE324208), CUT&RUN (GSE324820), and Xenium spatial transcriptomics (GSE324821).

## Abbreviations

C.APC: committed adipose progenitor cell
CNV: copy number variation
CTL: cytotoxic T lymphocytes
DDLPS: dedifferentiated liposarcoma
GEM: gel beads-in-emulsion
LPS: liposarcoma
MSC: mesenchymal stem cell
MSP: mesenchymal stem/progenitor
N-NMF: normalized non-negative matrix factorization
PCA: principal component analysis
Pre-ADC: pre-adipocyte
snATAC-seq: single-nucleus assay for transposase-accessible chromatin with sequencing
snMultiome: single nucleus multiome
snRNA-seq: single-nucleus ribonucleic acid sequencing
TAM: tissue-associated macrophage
Tem: Effector memory T cells
TF: transcription factor
TSS: transcription start site
UMAP: uniform manifold approximation and projection
UMI: unique molecular identifier
WD/DD: LPS well-differentiated/dedifferentiated liposarcoma
WDLPS: well-differentiated liposarcoma
WES: whole exome sequencing
WGS: whole genome sequencing

## ACKNOWLEDGMENTS

The authors acknowledge support from Sarcoma-Oma Foundation, the Moeller Foundation, the William Oats Foundation, and the MD Anderson Cancer Center Support Grant (P30 CA016672). The authors acknowledge the support of the High Performance Computing for research facility at the University of Texas MD Anderson Cancer Center for providing computational resources that have contributed to the research results reported in this paper. RAD is supported by NIH grant T32 CA009666 and an ASCO Young Investigator Award in honor of Grant and Victoria Merryman.

## REFERENCES

1. Mack, T. M. Sarcomas and other malignancies of soft tissue, retroperitoneum, peritoneum, pleura, heart, mediastinum, and spleen. Cancer 75, 211–244 (1995).

2. Thirasastr, P. & Somaiah, N. Overview of systemic therapy options in liposarcoma, with a focus on the activity of selinexor, a selective inhibitor of nuclear export in dedifferentiated liposarcoma. Ther Adv Med Oncol 14, 17588359221081073 (2022).

3. Lee, A. T. J., Thway, K., Huang, P. H. & Jones, R. L. Clinical and Molecular Spectrum of Liposarcoma. J Clin Oncol 36, 151–159 (2018).

4. Fletcher, C. D. et al. Correlation between clinicopathological features and karyotype in lipomatous tumors. A report of 178 cases from the Chromosomes and Morphology (CHAMP) Collaborative Study Group. Am J Pathol 148, 623–630 (1996).

5. Coindre, J.-M., Pédeutour, F. & Aurias, A. Well-differentiated and dedifferentiated liposarcomas. Virchows Arch 456, 167–179 (2010).

6. Knight, J. C., Renwick, P. J., Dal Cin, P., Van den Berghe, H. & Fletcher, C. D. Translocation t(12;16)(q13;p11) in myxoid liposarcoma and round cell liposarcoma: molecular and cytogenetic analysis. Cancer Res 55, 24–27 (1995).

7. Gronchi, A. et al. Variability in Patterns of Recurrence After Resection of Primary Retroperitoneal Sarcoma (RPS): A Report on 1007 Patients From the Multi-institutional Collaborative RPS Working Group. Ann Surg 263, 1002–1009 (2016).

8. Lahat, G. et al. Resectable well-differentiated versus dedifferentiated liposarcomas: two different diseases possibly requiring different treatment approaches. Ann Surg Oncol 15, 1585–1593 (2008).

9. Laurino, L., Furlanetto, A., Orvieto, E. & Dei Tos, A. P. Well-differentiated liposarcoma (atypical lipomatous tumors). Semin Diagn Pathol 18, 258–262 (2001).

10. Henricks, W. H., Chu, Y. C., Goldblum, J. R. & Weiss, S. W. Dedifferentiated liposarcoma: a clinicopathological analysis of 155 cases with a proposal for an expanded definition of dedifferentiation. Am J Surg Pathol 21, 271–281 (1997).

11. Dalal, K. M., Kattan, M. W., Antonescu, C. R., Brennan, M. F. & Singer, S. Subtype specific prognostic nomogram for patients with primary liposarcoma of the retroperitoneum, extremity, or trunk. Ann Surg 244, 381–391 (2006).

12. Singer, S., Antonescu, C. R., Riedel, E. & Brennan, M. F. Histologic subtype and margin of resection predict pattern of recurrence and survival for retroperitoneal liposarcoma. Ann Surg 238, 358–370; discussion 370-371 (2003).

13. Thway, K. Well-differentiated liposarcoma and dedifferentiated liposarcoma: An updated review. Semin Diagn Pathol 36, 112–121 (2019).

14. Horvai, A. E., DeVries, S., Roy, R., O’Donnell, R. J. & Waldman, F. Similarity in genetic alterations between paired well-differentiated and dedifferentiated components of dedifferentiated liposarcoma. Mod Pathol 22, 1477–1488 (2009).

15. Evans, H. L. Atypical lipomatous tumor, its variants, and its combined forms: a study of 61 cases, with a minimum follow-up of 10 years. Am J Surg Pathol 31, 1–14 (2007).

16. Chrisinger, J. S. A. et al. The degree of sclerosis is associated with prognosis in well-differentiated liposarcoma of the retroperitoneum. J Surg Oncol 120, 382–388 (2019).

17. Tap, W. D. et al. Evaluation of well-differentiated/de-differentiated liposarcomas by high-resolution oligonucleotide array-based comparative genomic hybridization. Genes Chromosomes Cancer 50, 95–112 (2011).

18. Beird, H. C. et al. Genomic profiling of dedifferentiated liposarcoma compared to matched well-differentiated liposarcoma reveals higher genomic complexity and a common origin. Cold Spring Harb Mol Case Stud 4, a002386 (2018).

19. Ozturk, N., Singh, I., Mehta, A., Braun, T. & Barreto, G. HMGA proteins as modulators of chromatin structure during transcriptional activation. Front Cell Dev Biol 2, 5 (2014).

20. Kanojia, D. et al. Genomic landscape of liposarcoma. Oncotarget 6, 42429–42444 (2015).

21. Keung, E. Z. et al. Increased H3K9me3 drives dedifferentiated phenotype via KLF6 repression in liposarcoma. J Clin Invest 125, 2965–2978 (2015).

22. Keung, E. Z. & Rai, K. H3K9me3-mediated repression of KLF6: Discovering a novel tumor suppressor in liposarcoma using a systematic epigenomic approach. Mol Cell Oncol 3, e1093691 (2016).

23. Pimenta, E. M. et al. Epigenetic dysregulation of metabolic programs mediates liposarcoma cell plasticity. Sci Transl Med 18, eadw4689 (2026).

24. Gruel, N. et al. Cellular origin and clonal evolution of human dedifferentiated liposarcoma. Nat Commun 15, 7941 (2024).

25. Merrick, D. et al. Identification of a mesenchymal progenitor cell hierarchy in adipose tissue. Science 364, eaav2501 (2019).

26. Cawthorn, W. P., Scheller, E. L. & MacDougald, O. A. Adipose tissue stem cells meet preadipocyte commitment: going back to the future. J Lipid Res 53, 227–246 (2012).

27. Maniyadath, B., Zhang, Q., Gupta, R. K. & Mandrup, S. Adipose tissue at single-cell resolution. Cell Metab 35, 386–413 (2023).

28. Rivera-Gonzalez, G. C. et al. Single-cell lineage tracing reveals hierarchy and mechanism of adipocyte precursor maturation. bioRxiv 2023.06.01.543318 (2023) doi:10.1101/2023.06.01.543318.

29. Truong, D. D. et al. Mapping the Single-cell Differentiation Landscape of Osteosarcoma. Clin Cancer Res 10.1158/1078-0432.CCR-24-0563 (2024) doi:10.1158/1078-0432.CCR-24-0563.

30. Terekhanova, N. V. et al. Epigenetic regulation during cancer transitions across 11 tumour types. Nature 623, 432–441 (2023).

31. Chen, Y., Xu, L., Lin, R. Y.-T., Müschen, M. & Koeffler, H. P. Core transcriptional regulatory circuitries in cancer. Oncogene 39, 6633–6646 (2020).

32. Matushansky, I. et al. A developmental model of sarcomagenesis defines a differentiation-based classification for liposarcomas. Am J Pathol 172, 1069–1080 (2008).

33. Pimenta, E. M. et al. Epigenetic dysregulation of metabolic programs mediates liposarcoma cell plasticity. bioRxiv 2025.01.20.633920 (2025) doi:10.1101/2025.01.20.633920.

34. McGinnis, C. S., Murrow, L. M. & Gartner, Z. J. DoubletFinder: Doublet Detection in Single-Cell RNA Sequencing Data Using Artificial Nearest Neighbors. Cell Syst 8, 329–337.e4 (2019).

35. Butler, A., Hoffman, P., Smibert, P., Papalexi, E. & Satija, R. Integrating single-cell transcriptomic data across different conditions, technologies, and species. Nat Biotechnol 36, 411–420 (2018).

36. Huang, M. et al. SAVER: gene expression recovery for single-cell RNA sequencing. Nat Methods 15, 539–542 (2018).

37. McInnes. UMAP: Uniform Manifold Approximation and Projection. Journal of Open Source Software 3, (2018).

38. Gulati, G. S. et al. Single-cell transcriptional diversity is a hallmark of developmental potential. Science 367, 405–411 (2020).

39. Cusanovich, D. A. et al. Multiplex single cell profiling of chromatin accessibility by combinatorial cellular indexing. Science 348, 910–914 (2015).

40. Granja, J. M. et al. ArchR is a scalable software package for integrative single-cell chromatin accessibility analysis. Nat Genet 53, 403–411 (2021).

41. Zhang, Y. et al. Model-based analysis of ChIP-Seq (MACS). Genome Biol 9, R137 (2008).

42. Skene, P. J., Henikoff, J. G. & Henikoff, S. Targeted in situ genome-wide profiling with high efficiency for low cell numbers. Nat Protoc 13, 1006–1019 (2018).

43. Skene, P. J. & Henikoff, S. An efficient targeted nuclease strategy for high-resolution mapping of DNA binding sites. Elife 6, e21856 (2017).

44. Li, H. et al. The Sequence Alignment/Map format and SAMtools. Bioinformatics 25, 2078–2079 (2009).

45. Ramírez, F. et al. deepTools2: a next generation web server for deep-sequencing data analysis. Nucleic Acids Res 44, W160–165 (2016).

46. Robinson, J. T. et al. Integrative genomics viewer. Nat Biotechnol 29, 24–26 (2011).

