## Supplemental Figures for "Spatially-resolved single cell atlas of liposarcoma reveals lineage hierarchies, immune niches, and regulatory circuits"

**SUPPLEMENTARY INFORMATION**

**Table S1. Characteristics of WD and DD LPS tumors included in the study.**

**Table S2. Annotation of samples.**

**Table S3. Clinical characteristics of LPS patients included in the study.**

**Table S4. Marker genes used for cluster annotation.**

**Table S5. CUT&RUN experimental parameters.**

**Table S6. Genes included in custom 480-gene Xenium spatial transcriptomic panel.**

**Supplementary Figure S1. Histologic validation of samples.**

Representative images of H&E staining of WDLPS and DDLPS specimens utilized for single nucleus multiome sequencing. There are some samples for which we analyzed tumor samples in duplicate, which is denoted with A and B (e.g. 2-WD-A and 2-WD-B). WDLPS tumors were classified as either adipocytic or sclerotic.

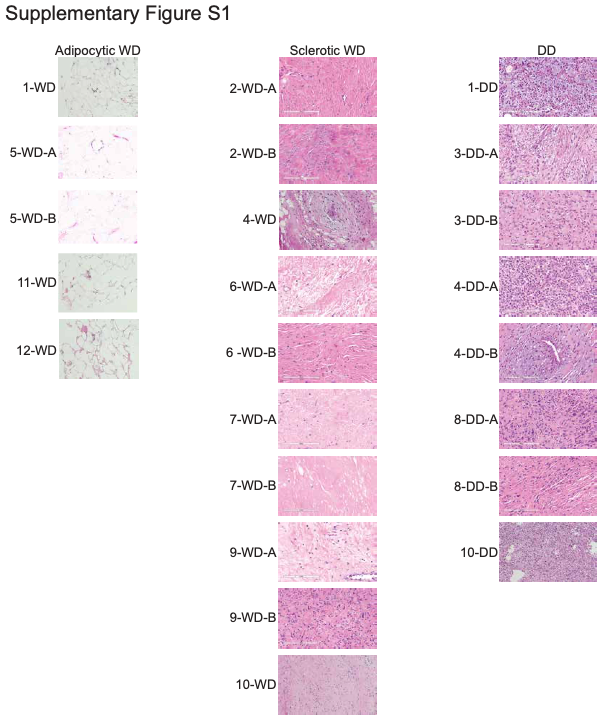

**Supplementary Figure S2. InferCNV analysis.**

(A) Infer CNV analysis showing copy number gains and loss in all chromosomes among all tumor clusters with benign cell clusters (4, 24, and 28) used for normalization. (B) Violin plot showing inferCNV scores by grouped clusters from snRNA-seq data. (C) Violin plot showing inferCNV scores by cluster from snRNA-seq data.

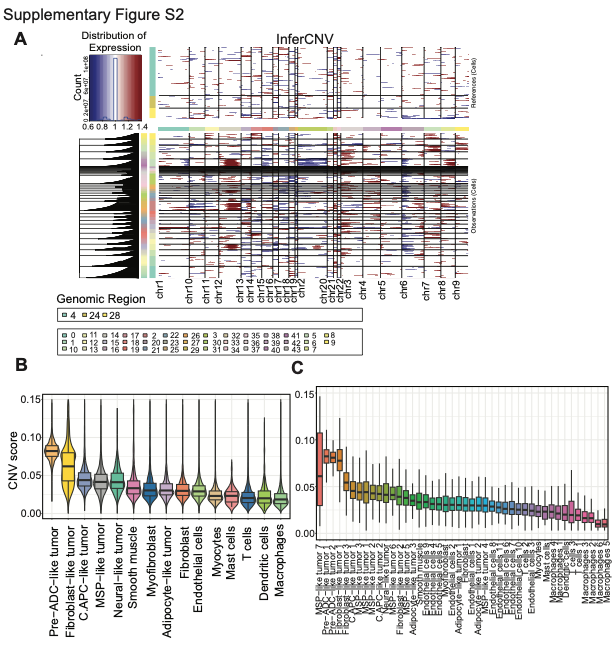

**Supplementary Figure S3. CytoTRACE analysis.**

(A) Violin plot showing CytoTRACE scores by cluster from snRNA-seq data. (B) Violin plot Violin plot showing CytoTRACE scores by grouped clusters from snRNA-seq data.

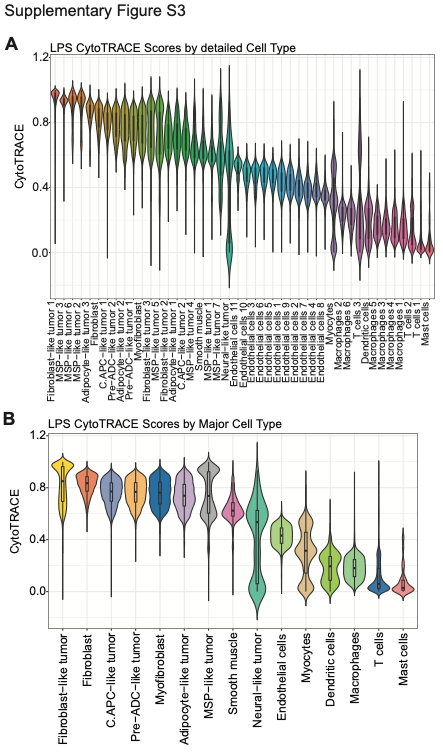

**Supplementary Figure S4. Individual cell types identified in snRNA-seq analysis.**

(A) UMAP visualization of individual groups of malignant cell clusters by type (e.g. MSP-like, C.APC-like, etc). (B) UMAP visualization of benign cell clusters by type (e.g. endothelial cells, macrophages, etc).

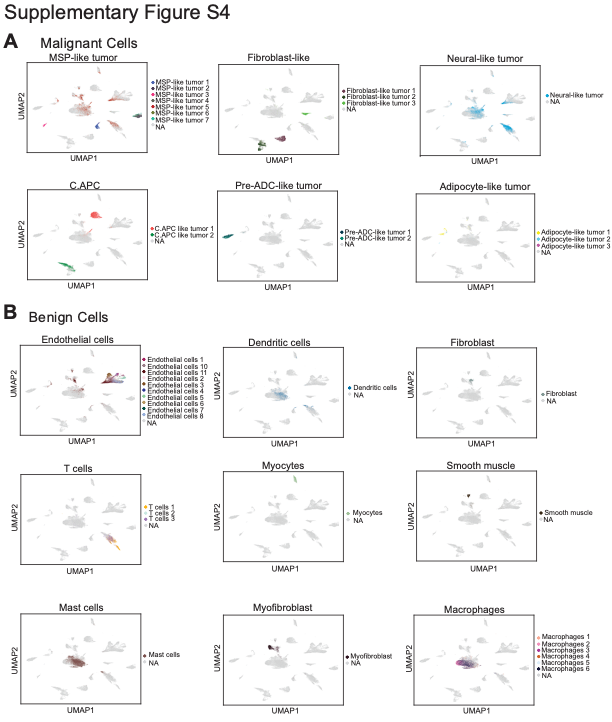

**Supplementary Figure S5. Archetype analysis.**

(A) Heatmap showing presence of archetypes within tumor cell clusters identified in snRNA-seq data. (B) UMAP analysis of archetypes with annotation based on cluster identify from snRNA-seq data. (C) UMAP analysis of archetypes with overlay of intensity of archetypes 3, 6, and 7. (D) Heatmap showing intensity of each archetype within each tumor cell cluster identified in snRNA-seq data. (E-P) Heatmaps showing genes whose expression best correlates with each archetype.

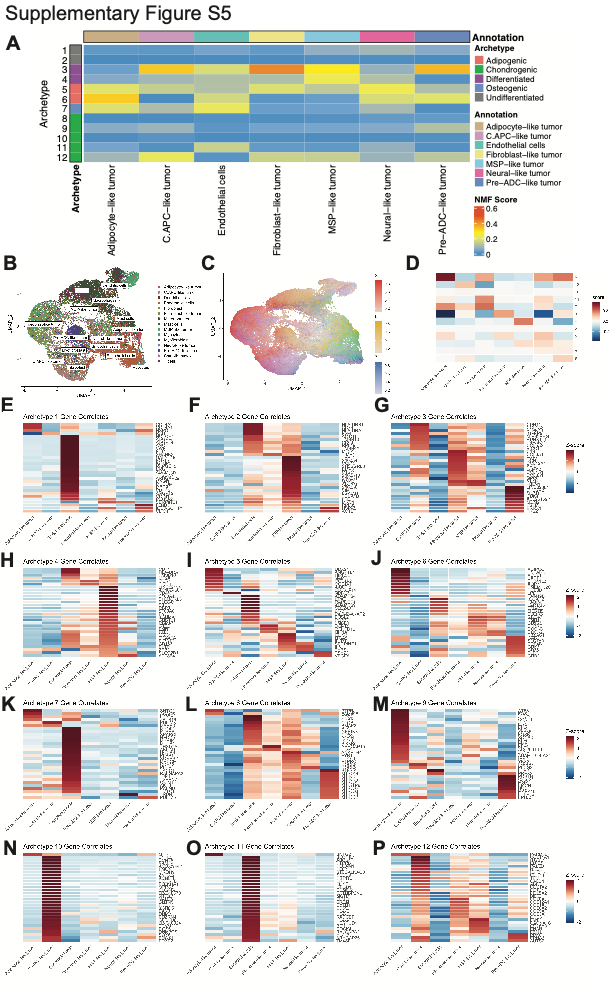

**Supplementary Figure S6. Distribution of cell types by tumor and subtype.**

(A) Stacked bar plot demonstrating proportion of each tumor cell cluster that is found in either adipocytic WD, sclerotic WD, or DD LPS. (B) Stacked bar plot demonstrating proportion of each combined tumor cell cluster that is found in either adipocytic WD, sclerotic WD, or DD LPS. (C) Number of cells of each cell cluster found in adipocytic WD, sclerotic WD, and DD LPS tumors. (D) Stacked bar plot demonstrating proportion of each tumor cell cluster represented by the individual tumors. (E) Proportion of each benign cell type that was found in either adipocytic WD, sclerotic WD, or DD LPS tumors.

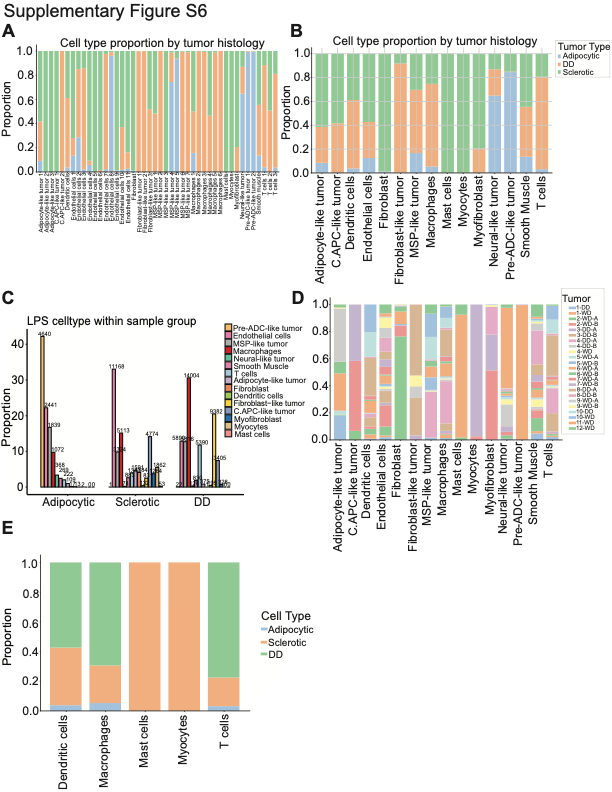

**Supplementary Figure S7. Comparison of single cell phenotypes in 3 matched WD/DD LPS pairs.**

(A) UMAP analysis of snRNA-seq data from the 3 matched pairs of WD/DD LPS tumors with each tumor specimen in a different color. (B) UMAP analysis demonstrating 36 unique clusters. (C) Heatmap showing marker genes that were used to annotate the clusters. (D) UMAP with MDM2 expression projected. Malignant WD/DD LPS cells are defined by MDM2 amplification. (E) Stacked bar plot showing the percent of cells in WD and DD tumors within each category.

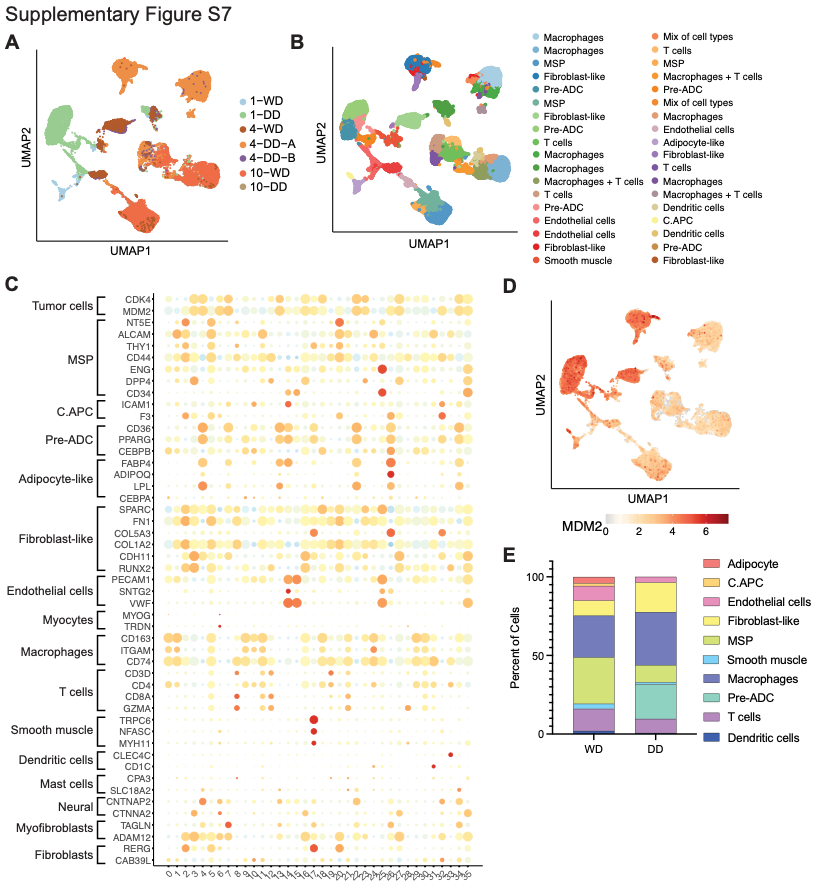

**Supplementary Figure S8. Annotation of Xenium spatial transcriptomic data.**

(A) Heatmap showing marker genes that were used to annotate clusters identified in Xenium data. (B) Representative UMAPs demonstrating expression of marker genes defining individual cell clusters, including FABP4 and ADIPOQ, (adipocyte-like), PPARG (C.APC), PDGFRA and NT5E (MSP), and CD24 (T cells).

**Supplementary Figure S9. Analysis of Xenium data in patient 10.**

(A-B) From left to right, post-analysis H&E-stained image, Xenium spatial transcriptomic maps, and spatial maps with individual adipocyte genes (ADIPOQ and FABP4) and mesenchymal genes (THY1 and KIT) in DD (A) versus WD (B). (C, F) Spatial map of niches in DD and WD components. (D, G) Niche composition of 5 niches identified. (E, H) Stacked bar plots showing proportion of cells by cluster type that made up each niche within DD and WD components.

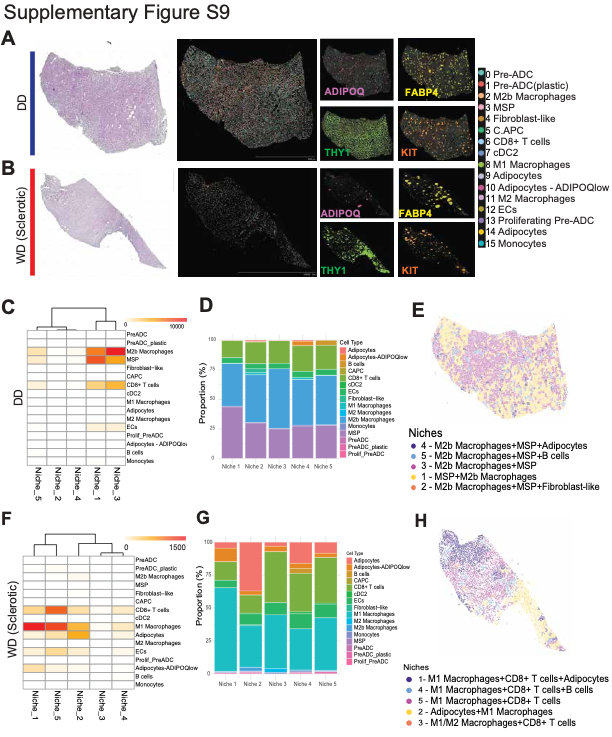

**Supplementary Figure S10. Analysis of Xenium data in patient 4.**

(A-B) From left to right, post-analysis H&E-stained image, Xenium spatial transcriptomic maps, and spatial maps with individual adipocyte genes (ADIPOQ and FABP4) and mesenchymal genes (THY1 and KIT) in DD (A) versus WD (B). (C, F) Spatial map of niches in DD and WD components. (D, G) Niche composition of 5 niches identified. (E, H) Stacked bar plots showing proportion of cells by cluster type that made up each niche within DD and WD components.

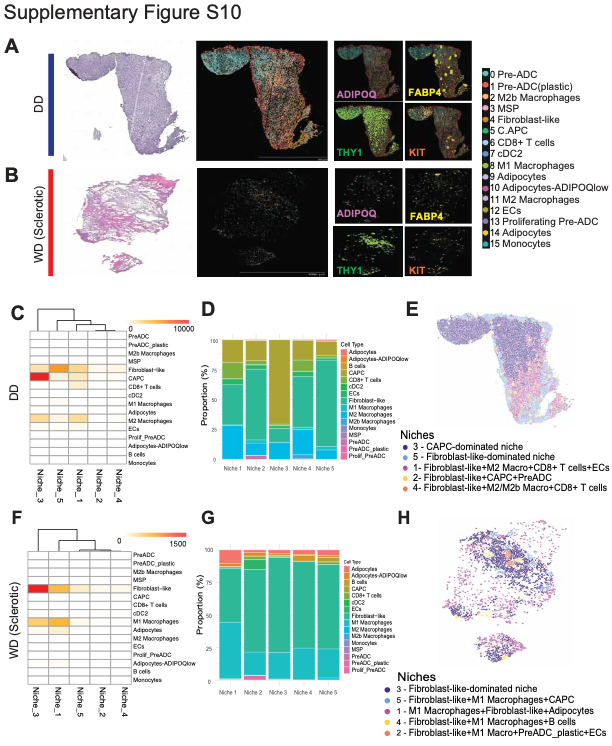

**Supplementary Figure S11. T cell and macrophage subsets.**

(A) Bubble plots of T cell markers by individual cluster (left), individual tumor (center), and tumor subtype (right). (B) Bubble plots of macrophage markers by individual cluster (left), individual tumor (center), and tumor subtype (right).

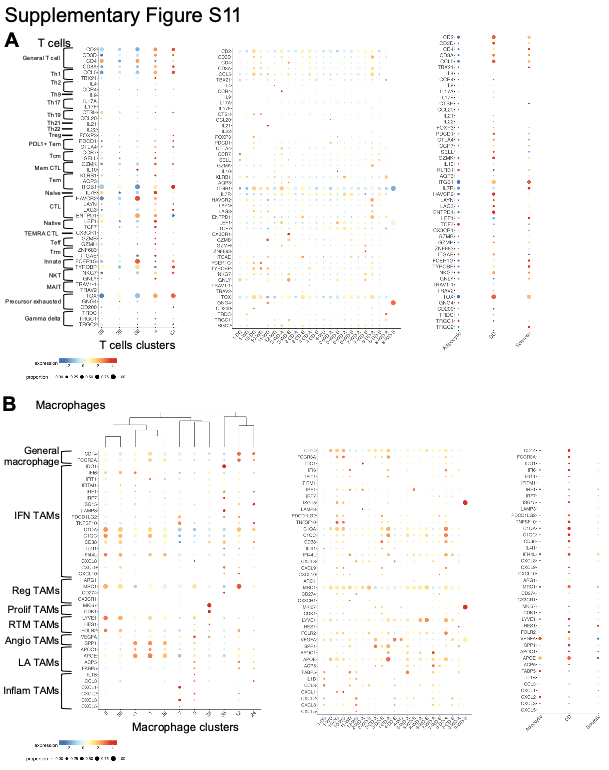

**Supplementary Figure S12. Assessment of chromatin openness in WD/DDLPS.**

(A) UMAP analysis of snATAC-seq data showing differences in profiles based on adipocytic WD, sclerotic WD, and DD. (B) UMAP analysis of snATAC-seq data with each color representing a unique tumor sample. (C) UMAP analysis of snATAC-seq data with each color representing a unique cluster. (D) UMAP analysis of snATAC-seq data from only the malignant cells (i.e. excluding immune cells) with each color representing a different cell subtype. (E) UMAP analysis of snATAC-seq data from only malignant cells with each color representing a different cell cluster.

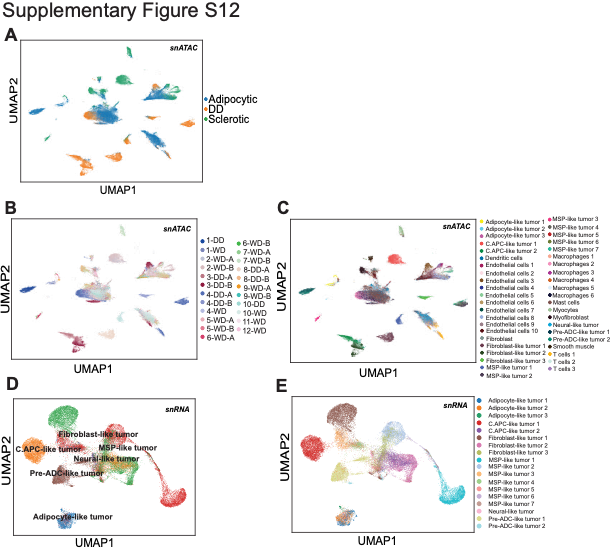

**Supplementary Figure S13. ATAC peak enrichment.**

(A-B) Heat maps showing peaks enriched in snATAC-seq data by individual cluster (A) and tumor sample (B).

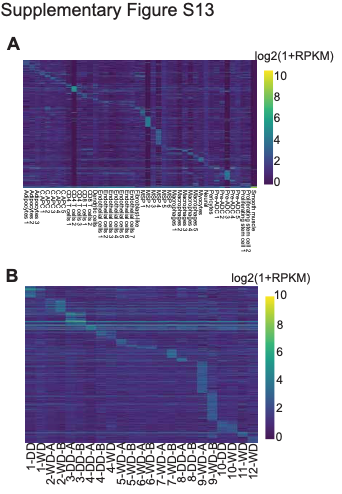

**Supplementary Figure S14. TF motif enrichment by cluster.**

(A-B) Motif analysis using Homer identified motifs enriched in malignant cell clusters (A) and benign cell clusters (B).

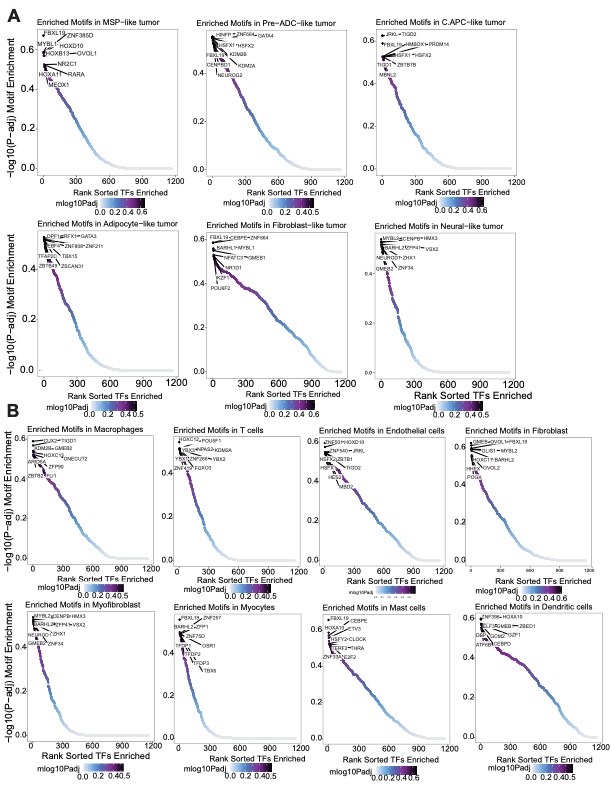

**Supplementary Figure S15. Analysis of H3K4m3 and H3K27me3 CUT&RUN.**

(A) PCA analysis of H3K4me3 CUT&RUN data. (B) Correlation heatmap analysis between biological replicates. (C) Differential peak analysis identified 3630 H3K4me3 peaks enriched in WDLPS and 291 peaks enriched in DDLPS. (D) Integrative genome browser view of CUT and RUN analysis showing H3K4me3 enrichment at the PPARG locus in WD and the KIT locus in DD. (E) PCA analysis of H3K27me3 CUT&RUN data. (F) Correlation heatmap analysis between biological replicates. (G) Differential peak analysis identified 3630 H3K27me3 peaks enriched in WDLPS and 291 peaks enriched in DDLPS. (H) Integrative genome browser view of CUT and RUN analysis showing H3K27me3 enrichment at the CD6 locus in WD and the OSBPL8 locus in DD.

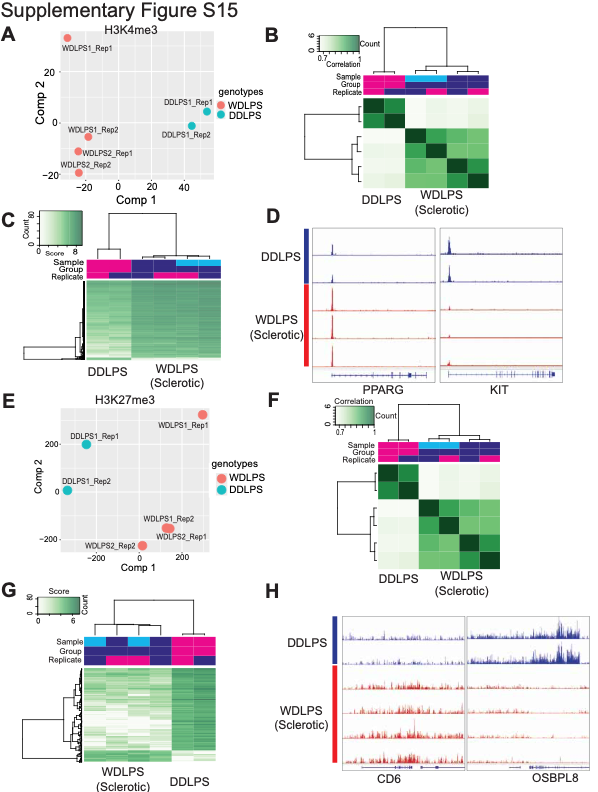

**Supplementary Figure S16. Identification of core TF circuits in LPS**

(A) Workflow integrating snATAC-seq, snRNA-seq, and H3K27ac CUT&RUN to define CRC-like TFs. Top 50 TF motifs enriched in tumor snATAC clusters from adipocytic and sclerotic WDLPS and DDLPS were screened for concordant TF and target gene expression and enhancer gains at their regulatory loci. (B) Expression of top 50 TFs ranked by motif enrichments in adipocytic and sclerotic WDLPS and DDLPS. (C) Expression of predicted direct targets of CRC-candidate TFs, showing enrichment of proliferative and anti-adipogenic programs in sclerotic WDLPS and activation of EMT and osteo/chondrogenic pathways alongside reduced adipogenic maturation in DDLPS.

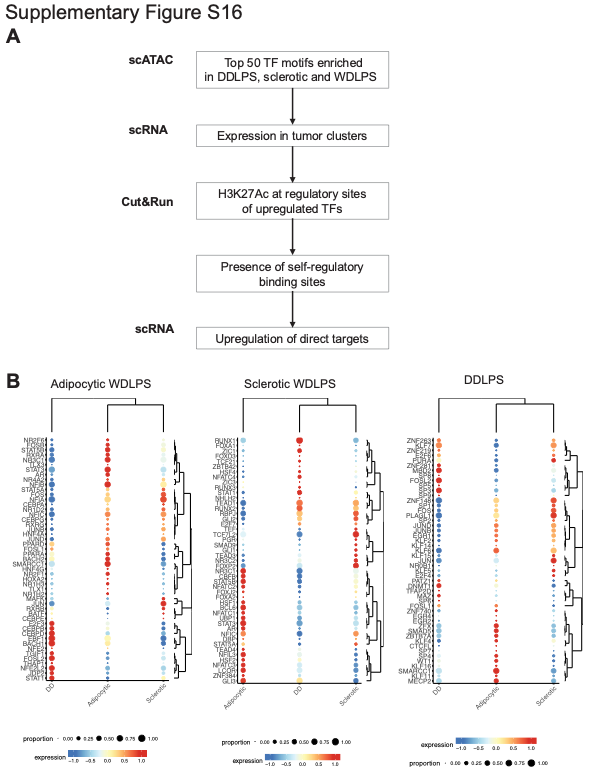

**Supplementary Figure S17. Determination of self-regulating enhancers in WD and DD LPS**

(A) Integrative genome browser view of TFs meeting criteria of motif enrichment, elevated expression, and CUT&RUN H3K27ac enhancer gains with self-regulatory binding sites in sclerotic WDLPS and DDLPS are shown. CRC-candidate TFs identified include GLI2, TCF7L2, and RBPJ (sclerotic WDLPS) and KLF7, FOSL2, and SP3 (DDLPS). (B) Bubble plot demonstrating gene expression of the core CRC-candidate TFs and their downstream targets in adipocytic and sclerotic WD and DD tumor cells. For GLI1, targets are PTCH1, CCND1, PDGFRB, and CDKN2.

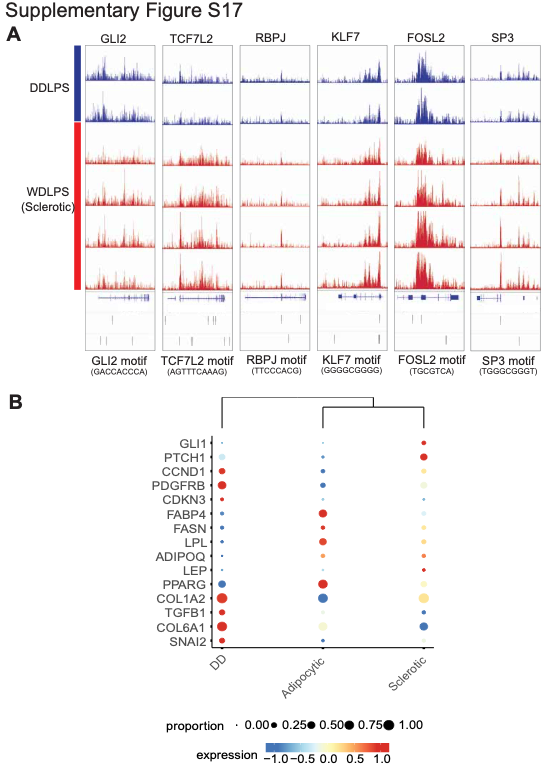
